# Modeling tactile pleasantness across skin types at the individual level reveals a reliable and stable basic function

**DOI:** 10.1101/2022.04.11.487838

**Authors:** Laura Crucianelli, Marie Chancel, H. Henrik Ehrsson

## Abstract

Touch is perceived most pleasant when delivered at velocities known to optimally activate C Tactile afferents. At the group level, pleasantness ratings of touch delivered at velocities in the range 0.3-30 cm/s follows an inverted-U shape curve, with maximum pleasantness between 1 and 10 cm/s. However, the prevalence, reliability, and stability of this function at the individual level and across skin types remains unknown. Here, we tested a range of seven velocities delivered with a soft brush, on both hairy and non-hairy skin in 123 participants. We showed that the relationship between pleasantness and velocity of touch is significantly best described by a negative quadratic model at the individual level in the majority of participants both on hairy (67.1%) and non-hairy skin (62.6%). Higher interoceptive accuracy and self-reported depression were related to a better fit of the quadratic model and to the steepness of the curve, respectively. The prevalence of the quadratic model was stable across body sites (62.6%), across two experimental sessions (73-78%,), and regardless of the number of trials. Thus, the individual perception of tactile pleasantness follows a characteristic velocity-dependent function across skin types and shows trait characteristics, making it a possible biomarker for mental health disorders.

## Introduction

Research on the sense of touch has identified two distinct modalities; a discriminative and an affective one (see McGlone, Wessberg and Olausson, 2014, for a review). The sensory-discriminative dimension supports the detection and identification of external stimuli, providing information about physical characteristics and spatial location. In contrast, the motivational-affective dimension is involved in the valence and motivational nature of the stimuli, such as hedonic and emotional relevance (Olausson et al., 2002; McGlone et al., 2007; Olausson et al., 2010; Löken et al., 2009). The affective dimension of touch can be investigated by means of a low-pressure, slow, caress-like tactile stimulation delivered at velocities between 1 and 10 cm/s (Löken et al., 2009). This type of touch is perceived as more pleasant compared to slower or faster velocities according to subjective ratings. Studies using microneurography, a neurophysiological method allowing the recording of the activity of single peripheral nerves in the skin (Ackerley & Watkins, 2018), showed an activation of C-tactile (CT) afferents when touch presents the aforementioned characteristics (Vallbo et al., 1999).

Over the last few decades, research has supported the hypothesis that CT afferents constitute a distinct system, both from the anatomical and functional point of view, which may contribute to responses to slow, caress-like types of touch and provide pleasant sensations (Vallbo et al., 1999; Olausson et al., 2002; Löken et al., 2009). These fibers are mainly present in the hairy skin of the body (Vallbo et al., 1999; Watkins, Dione, Ackerley, Backlund Wasling, Wessberg & Löken, 2021), and when these fibers are activated, individuals report a pleasant percept. Löken and colleagues showed that there was a linear correlation between the activation of CT fibers and the subjective report of pleasantness (Löken et al., 2009). At the group level, the averaged pleasantness ratings of touch delivered at velocities in the range between 0.3 and 30 cm/s follow an inverted-U shape curve, with maximum pleasantness between 1 and 10 cm/s (Löken et al., 2009). This pattern of relationship between velocity of touch and tactile pleasantness has been replicated at the group level across several studies (see Cruciani et al., 2021 for a recent meta-analysis). However, perception occurs in individuals, not groups, and we do not know how prevalent the inverted-U shape curve is and if it observable in most people as the prevailing CT-hypothesis would predict. One concern is that non-linear effects found at the group level could be driven by a minority of participants or effects that are so small as being psychologically “meaningless” for individuals (i.e., only an aggregated group-level effect).

To the best of our knowledge, only one recent study has investigated the fitting of the inverted U-shape curve at the individual level on hairy skin (Croy, Bierling, Sailer & Ackerley, 2021). Croy et al. (2021) pooled data from 5 separate studies and showed that surprisingly only 42% of participants presented the typical inverted U-shape curve, while 44% were better described by a linear model, and the remaining participants showed no significant effect of the velocity of touch on pleasantness ratings. Moreover, from the study of Croy and colleagues we do not know if the shape of the inverted U-function varies across different skin types with different CT innervation within individuals. CT afferents are seven times more numerous on hairy (forearm) compared to non-hairy skin (i.e., palm, Watkins et al., 2021) and, according to a central prediction of the CT-hypothesis, this difference should be reflected in more prevalent inverted-U shape pleasantness curves on hairy skin, but this important question has not been examined at the individual level. Noteworthy, and a recent meta-analysis at the group level found no significant differences between hairy and non-hairy skin in the perception of tactile pleasantness (Cruciani, Zanini, Russo, Boccardi, & Spitoni, 2021), although the inverted-U shape was not analyzed at the individual subject level. Nevertheless, such evidence questions the traditional distinction between hairy and non-hairy skin in the perception of affective touch and what is known so far of the role that the CT system plays is mediating gentle touch, whereby the palm of the hand has classically been used as a control body site to compare against the forearm. Thus, we believe that now more than ever there is the need of more studies that systematically investigate the differences in tactile pleasantness across hairy and non-hairy skin sites. Accordingly, here we aimed to investigate the relationship between velocity of touch and pleasantness scores at the individual level, characterizing individual differences and prevalence of inverted U-shape functions across hairy and non-hairy skin as well as testing the stability and reliability of such individual pleasure-velocity functions across sessions and trials repetitions.

Importantly, the specialized peripheral and central neurophysiological systems underpinning the perception of affective touch (including the CT-system) seem to take a different pathway than the discriminative, more emotionally neutral touch from the peripheral nerves of the skin to the posterior insular cortex (Olausson et al., 2002, but see Gazzola et al., 2012). A recent study has suggested that spinothalamic projections might play only a partial role in the hedonic processing of touch mediated by C-tactile afferents, thus proposing a more integrated view of tactile pleasantness (Marshall et al., 2019). However, functional imaging studies in humans agree on the idea that the posterior insular cortex as a primary cortical target for C fibers (Olausson et al., 2002; Morrison, 2016), an area strongly interconnected with regions related to emotional processing such as the amygdala, hypothalamus, and orbital frontal cortex. In light of these neurophysiological observations, affective touch has been reconceptualized as an interoceptive modality providing information about the internal states of the body (Craig, 2002; 2009; Björnsdotter, Morrison, & Olausson, 2010; Crucianelli et al., 2018).

In parallel with the growing understanding of the functions, pathways, and characteristics of the affective touch system, there has also been increasing interest in better understanding the implications of dysfunction in such a system. In particular, a few studies have investigated the perception of affective touch in clinical populations, showing significant disruptions in the perception of affective touch in anorexia nervosa (Crucianelli et al., 2016), autism (Kaiser et al., 2016), right-hemisphere stroke (Kirsch et al., 2020), and chronic pain (Case et al., 2016), among others. Overall, these studies have suggested the existence of a link between mental health and social touch perception (Croy et al., 2016). Given these conceptualizations, it is important to understand whether the perception of affective touch could be used as a behavioral biomarker for the early identification of people at risk of developing a mental health disorder. For this approach to work, it is essential to clarify the reliability and stability of the affective touch modeling results in at the individual level in healthy samples.

The subjective perception of affective touch is the result of a combination of peripheral activation of skin receptors and central processing of such stimulation (Sailer, Hausmann & Croy, 2020). Furthermore, personality and psychological traits can also play a role in the way we perceive touch. Although touch is mediated by the skin, its perception is the result of a combination of bottom-up signals (i.e., afferent signals resulting from the activation of the nerve fibers in the skin) and top-down factors, such as previous experiences, the context, the identity and gender of the toucher (Ellingsen, Leknes, Løseth, Wessberg & Olausson, 2016). Accordingly, an additional aim of this study was to investigate the relationship between individual traits and characteristics (i.e., depression, anxiety, eating disorders symptomatology) and individual differences in the perception of tactile pleasantness. In line with the idea of affective touch as an interoceptive modality (Craig, 2002), we also aimed to investigate whether cardiac interoceptive accuracy, measured by means of a classic heartbeat-counting task (Schandry, 1981), would predict the pattern of the relationship between velocity of touch and tactile pleasantness (see Crucianelli et al., 2018 for a similar approach).

Here, across three experiments, we aimed to assess 1) whether and to what extent the classic negative quadratic model (i.e., inverted U shape) outperformed a linear model to describe the relationship between velocity and pleasantness of touch at the individual level; 2) which individual difference factors can predict the significance and prevalence of the negative quadratic model; 3) whether the relationship between velocity and pleasantness varies across hairy vs. non-hairy skin within individuals; and 4) whether the individual variability in the perception of tactile pleasantness is temporally stable, both when tested across two identical experimental sessions one week apart and when tested with an increasing number of repetitions at each velocity. We separately describe the specific procedures for each experiment below.

## EXPERIMENT 1

The *first aim* of Experiment 1 was to test whether the impact of stroking velocity on subjective pleasantness could be better described by a quadratic or a linear regression. Croy, Bierling, Sailer & Ackerley (2021) recently showed that the typical inverted U-shape curve described the relationship between the velocity of touch and pleasantness in only 42% of participants. However, these results were obtained by pooling data from 5 separate studies that included a total of 127 participants tested by different experimenters and under different conditions. Indeed, Croy and colleagues (2021) reported a consistent effect of the experimental setting on tactile pleasantness. Here, we aimed to investigate whether the classic negative quadratic model outperformed a linear model in describing the relationship between velocity and pleasantness at the individual level with a set of participants tested under the same conditions and by the same experimenter. We hypothesized that the majority of individual participants should display a better fit with negative quadratic model in line with the CT-hypothesis.

Furthermore, several factors are known to influence the shape of the group-level curve. Thus, the *second aim* of Experiment 1 was to test whether these individual characteristics can also modulate pleasantness ratings at the individual level. We collected data regarding several individual characteristics that could potentially influence the perceived pleasantness of touch based on previous studies. We included the Eating Disorder Examination Questionnaire (EDE-Q) to target potential body dissatisfaction and weight restraint behaviors (see Crucianelli et al., 2016; 2021); self-reported measures of depression and anxiety to target potential links with anhedonia and affective disorders symptomatology (Croy et al., 2016; Triscoli, Croy & Sailer, 2019; Lapp & Croy, 2021); and cardiac interoceptive accuracy, measured by means of the classic heartbeat-counting task (Schandry, 1981), to target interoceptive abilities quantified via a non-tactile channel. In keeping with the aforementioned literature, participants with higher scores on the EDE-Q, depression, and anxiety scales were expected to give lower pleasantness ratings overall and to have difficulties differentiating tactile pleasantness levels based on stroking velocities. In other words, we predicted that higher EDE-Q, depression, and anxiety scores would lead to “flatter curves”, i.e., the participants would give lower pleasantness ratings and would rate stroking at different velocities as being more similar. Moreover, because of this expected loss of influence of stroking velocity on pleasantness ratings, the quadratic model should be less relevant in participants with higher scores on these individual characteristics with respect to a random pattern of answers. In contrast, a higher cardiac interoceptive accuracy should be related to an increased perception of pleasantness of CT-optimal velocities (see also Crucianelli et al., 2018). Finally, we explored whether the same individual characteristics could predict not only how well participant’s answers would be described by a quadratic model but also how well the estimated coefficients of this quadratic model fit.

The *third aim* of Experiment 1 was to assess the hypothesized differences in velocity-pleasantness functions across hairy and non-hairy body sites (forearm/hairy skin vs. palm/non-hairy skin) based on the well-known differences in CT afferents density and related tactile pleasantness perception between these two skin types (e.g., Vallbo et al., 1999; McGlone et al., 2012; Watkins, Dione, Ackerley, Backlund Wasling, Wessberg & Löken, 2021). The CT-hypothesis predicts a more pronounced inverted U-shape pleasantness-velocity function on hairy skin and that a higher proportion of participants should express this function on hairy skin. To this end, we compared the observed and predicted likelihood for each participant to best fit negative quadratic model or the linear model for both body sites.

## Methods and Materials

### Participants

A total of 107 healthy, naïve participants (53 women and 54 men) were recruited for Experiment 1 using social media and advertising on the Karolinksa Institutet campus. An a priori power analysis based on previous studies in the field of affective touch (e.g., Croy, Bierling, Sailer & Ackerley, 2021) suggested that the present sample provided enough power to detect our effects of interest. The inclusion criteria included being 18-40 years old (26.2 ± 5 years) and being right-handed. The exclusion criteria included having a history of any psychiatric or neurological conditions, taking any medications, having sensory or health conditions that might result in a skin condition (e.g., psoriasis), and having any scars or tattoos on their left forearm or hand. The study was approved by the Swedish Ethical Review Authority. All participants provided signed written consent, and they received a cinema ticket as compensation for their time. The study was conducted in accordance with the provisions of the Declaration of Helsinki 1975, as revised in 2008. Part of the data obtained in Experiment 1 was analyzed using a different statistical method for different purposes and has been published as part of another manuscript (Crucianelli, Enmalm & Ehrsson, 2021).

### Self-reported measures

Participants were asked to provide demographic information, such as age and handedness. Next, the participants were asked to complete the following self-report questionnaires: the Eating Disorders Examination Questionnaire (EDE-Q 6.0), a 28-item questionnaire measuring eating disorders symptomatology that has good consistency and reliability (global score α = .90; Fairburn & Beglin, 1994, 2008; Peterson et al., 2007), and the Depression, Anxiety and Stress Scale–21 Item (DASS), a 21-item, three-scale self-reported measure of depression, anxiety and stress that has an α >.88. (Lovibond and Lovibond, 1995; Henry & Crawford, 2005).

### Cardiac interoceptive accuracy: heartbeat counting task (HCT)

The task is fully described in Appendix 1.

### Affective touch task

This task takes advantage of the discovery that affective, hedonic touch on the skin can be reliably elicited by soft, light stroking at specific velocities within the range of 1-10 cm/s that activate a specialized peripheral system of CT afferents (Löken, Wessberg, McGlone & Olausson, 2009; McGlone, Vallbo, Olausson, Löken, & Wessberg, 2007). Before starting the task, the experimenter identified two adjacent areas measuring 9×4 cm on the left forearm and two on the left palm with a washable marker (see Procedure for more details). Then, participants were familiarized with the rating scale ranging from 0, not at all pleasant, to 100, extremely pleasant. The touch was delivered using a soft brush (i.e., precision cheek brush No 032, Åhléns, Sweden) on the left forearm (hairy skin that contains CT afferents) and left palm (non-hairy skin, where only limited CT afferents activity has been reported; Watkins et al., 2021), and the task of the participants was always to verbally rate the pleasantness of the touch using the rating scale. Participants were asked to wear a blindfold so that they could better focus on the tactile experience. The touch was delivered at seven velocities (0.3, 1, 3, 6, 9, 18 and 27 cm/s). The two slow velocities of 3 and 6 cm/s are typically perceived as more pleasant (i.e., CT optimal velocities) than the borderline optimal velocities (1 and 9 cm/s) and the CT non-optimal speeds (0.3, 18 and 27 cm/s; Löken et al., 2009). In keeping with previous studies (e.g., Croy et al., 2021), each velocity was presented three times, for a total of 21 stroking trials per location (palm and forearm, in randomized order).

### Experimental Procedure

Participants were welcomed in the experimental room, and they were asked to sit at a table opposite the experimenter. Upon arrival, they were asked to sign the consent form and to complete the questionnaires presented in an online format: demographic questionnaire, EDE-Q and DASS. The questionnaires were always presented at the beginning of the experimental procedure to ensure that participants were given some time to rest before completing the heartbeat counting task, which was the first interceptive task that all participants completed. Given that previous studies showed that the heartbeat counting task might be influenced by other activities (Ring et al., 2015; Brener and Ring, 2016), we decided to keep it as the first task. Participants were given the choice to keep their eyes closed or open, allowing them to feel more comfortable and be as accurate as possible. Next, participants were asked to wear a disposable blindfold to complete the affective touch task. Participants were familiarized with the pleasantness rating scale, and the experimenter identified and marked with a washable marker two identical areas of 9×4 cm on the left forearm and palm, as in previous studies (Crucianelli et al., 2013; 2016; 2018). This was done to control for the stimulated area and for the pressure applied during the touch by checking that the tactile stimulation was only applied inside the marked areas (more pressure would result in a wider spreading of the brush, that is, the tactile stimulation would be applied outside of the marked borders). Alternating the stimulated areas prevented fatigue of the CT fibers (McGlone et al. 2012).

### Design and plan of analysis

To test our *first aim* (i.e., whether the impact of stroking velocity on tactile pleasantness was better described by a negative quadratic or a linear regression), the data acquired from the different body sites were considered separately. Please note that in the rest of the manuscript, we talk about quadratic model for simplicity, but we have always considered a *negative* quadratic function. For each participant, both types of regressions were performed on the mean ratings for each stroking velocity; the fitting procedures were performed in R. We first considered how many participants showed a significant main effect of velocity on the pleasantness ratings (ANOVA). Those who did not, were categorized as having a ‘random’ profile (i.e., not significantly described by either a quadratic or linear regression). The remaining participants were categorized either as having a ‘quadratic’ or ‘linear’ profile. To do so, we fitted two models: one linear: pleasantness ∼ velocity, and one quadratic: pleasantness ∼ velocity + velocity^2^. This categorization was straightforward when only the quadratic fit or the linear fit were significant. When both fits were significant, a likelihood-ratio test (LRT) was performed: this LRT compares the goodness-of-fit of our two models (quadratic versus linear) while considering the lesser parsimony of the quadratic model, i.e., the quadratic model includes one more parameter than the linear model and thus has an inherent advantage. A significant LRT (p < 0.05) meant the participants’ profile would be categorized as ‘quadratic’, otherwise the participant’s profile would be categorized as ‘linear’. We further quantified the goodness-of-fit of the models using the root mean square error (RMSE). A lower RMSE indicated a better fit.

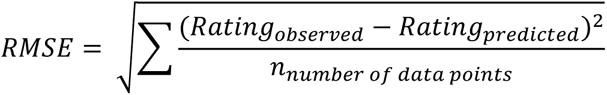

Moreover, to compare the relative relevance of each model at the individual level for our whole sample, we performed a confidence interval analysis (CI analysis) on the residual standard deviation (RSD). Simply put, in addition to knowing which model fits best, we wanted to know how much better one model was compared to the other for each participant and whether this difference in performance between the two models was meaningful at the group level. The RSD reflects how much the observed data spread around the regression curve and takes into account the degree of freedom of each model. The smaller the residual standard deviation was, the better the fit and the more predictive the model was.

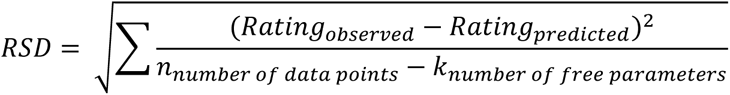

To compare the two models, the difference in RSD between the quadratic and linear models was calculated and summed across participants. We estimated a confidence interval using bootstrapping: 107 random RSD differences (quadratic – linear) were drawn with replacement from the actual participant’s RSD differences and summed; this procedure was repeated 10000 times to compute the 95% CI. Negative CI bounds would provide evidence of an overall better quadratic fit compared to a linear fit (see also Appendix Figure 1 in Appendix 1)

To address the *second aim* (i.e., to test whether individual characteristics influenced the pleasantness ratings at the individual level), we tested two correlation hypotheses: first, whether the quadratic RMSE increased with higher EDE-Q, depression, and anxiety scores, i.e., we expected a worst fit by the quadratic model. Second, whether the difference in RSD between the quadratic and the linear model (RSDd) would also increase, i.e., we assessed whether the relative relevance of the quadratic model compared to the linear model was reduced. Inverse correlations were expected with cardiac interoceptive accuracy: the lower the cardiac interoceptive accuracy was, the flatter the curve.

Furthermore, we tested the hypothesis that the same individual characteristics could predict not only how well participants’ answers would be described by a quadratic model but also the coefficients of this quadratic fit. Indeed, as mentioned above, the participants with higher EDE-Q, depression, and anxiety scores were expected to show “flatter curves”, i.e., to give lower pleasantness ratings and to rate the different stroking velocities as similarly pleasant. For the participants with a significant quadratic fit, the quadratic term coefficient (A2) reflects how different the pleasantness ratings are from one velocity to the other, i.e., the curvature of the curve. The intercept coefficient (A0) reflects the overall pleasantness (see Appendix 1 and Appendix Figure 2 for more details about the influence of each coefficient of the quadratic model on the collected pleasantness ratings). Based on our hypothesis, for participants with higher EDE-Q, depression, and anxiety scores, A2 should get closer to 0 and A0 should decrease. We tested these hypotheses by means of one-sided Spearman correlation tests on our whole population to investigate the relationship between the individual characteristics and the quadratic model RMSE and RSDd values. Then, in the subpopulation for which the quadratic fit was significant, we used one-sided Spearman correlation tests between the individual characteristics and the A2 (quadratic term) and A0 (intercept) coefficients.

The *third and last aim* of Experiment 1 was to assess the stability of the relationship between pleasantness ratings and stroking velocity across body sites (forearm versus palm). We compared the observed and predicted likelihood for a participant to be in the same profile category for both body sites. The predicted likelihood was calculated as the square of the probability of having a given profile for stroking on the forearm. The observed likelihood was calculated as the product of the probability of having a given profile for stroking on the forearm multiplied by the probability of participants with this given profile also having this profile for stroking on the palm. This analysis is similar to the one used by Bendas et al. (2021) to assess the stability of pleasantness ratings across repetitions on two different days. Finally, the participants who showed the ‘quadratic’ profile in both body sites could still display very different patterns of response (e.g., steep versus flat curves, different maximum and minimum). Thus, we investigated how correlated the coefficients of the quadratic model, A2, A1, and A0 were between the forearm and the palm stroking sites.

## Results

### Demographic and self-reported data

The mean scores and standard deviations for the EDE-Q and DASS are reported in Table S1. No effect of gender on any of these measures was found, with the exception of the EDE-Q scores. See also Appendix Figure 3 in Appendix 1 for the data distribution of the self-reported scores.

### Pleasantness for different stroking velocities on the forearm (hairy skin)

#### a. Categorization into ‘quadratic’, ‘linear’, or ‘random’ profiles

At the group level, we observed the classic inverted U-shaped curve. These results replicating the classically observed group-level results for tactile pleasantness ratings can be found in the Appendix 1 (see Appendix Figures 4 and 5). At the individual level, 25 participants (23.4%) did not show a significant effect of velocity on perceived pleasantness and were thus categorized as ‘random’. Out of the 82 remaining participants, 72 (67.1%) showed a ‘quadratic’ profile, and 10 (9.3%) showed a ‘linear’ profile (Figure 1). The raw sum for the RSD differences between the quadratic and linear models was - 620, and the 95% CI analysis confirmed that the quadratic regression outperformed the linear regression (lower bound: -727; upper bound: -516). Overall, our results showed that despite individual differences, the influence of stroking velocity on pleasantness ratings was better described by a quadratic regression than by a linear regression at the individual level as it was at the group level.

**Figure 1:**
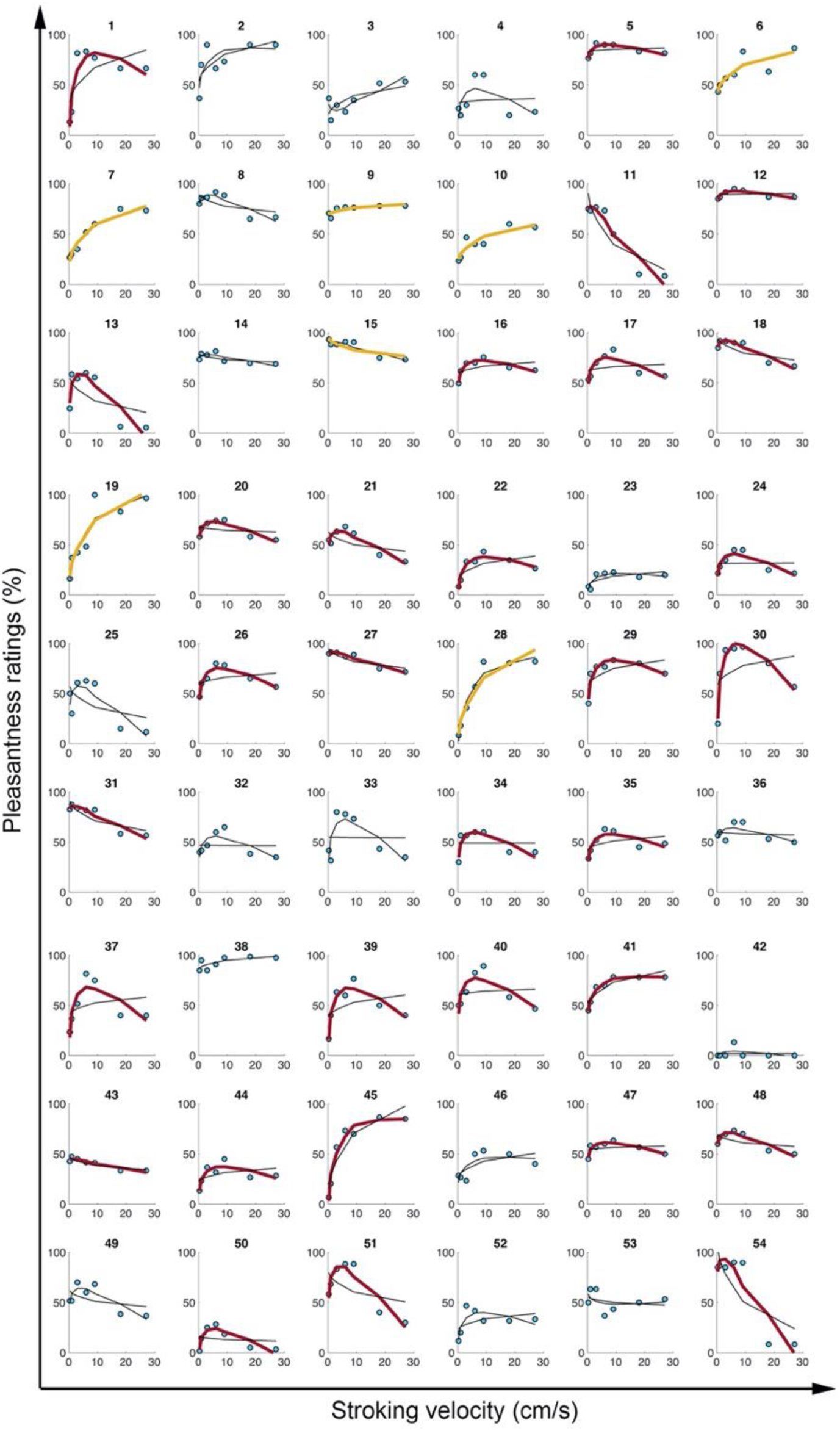
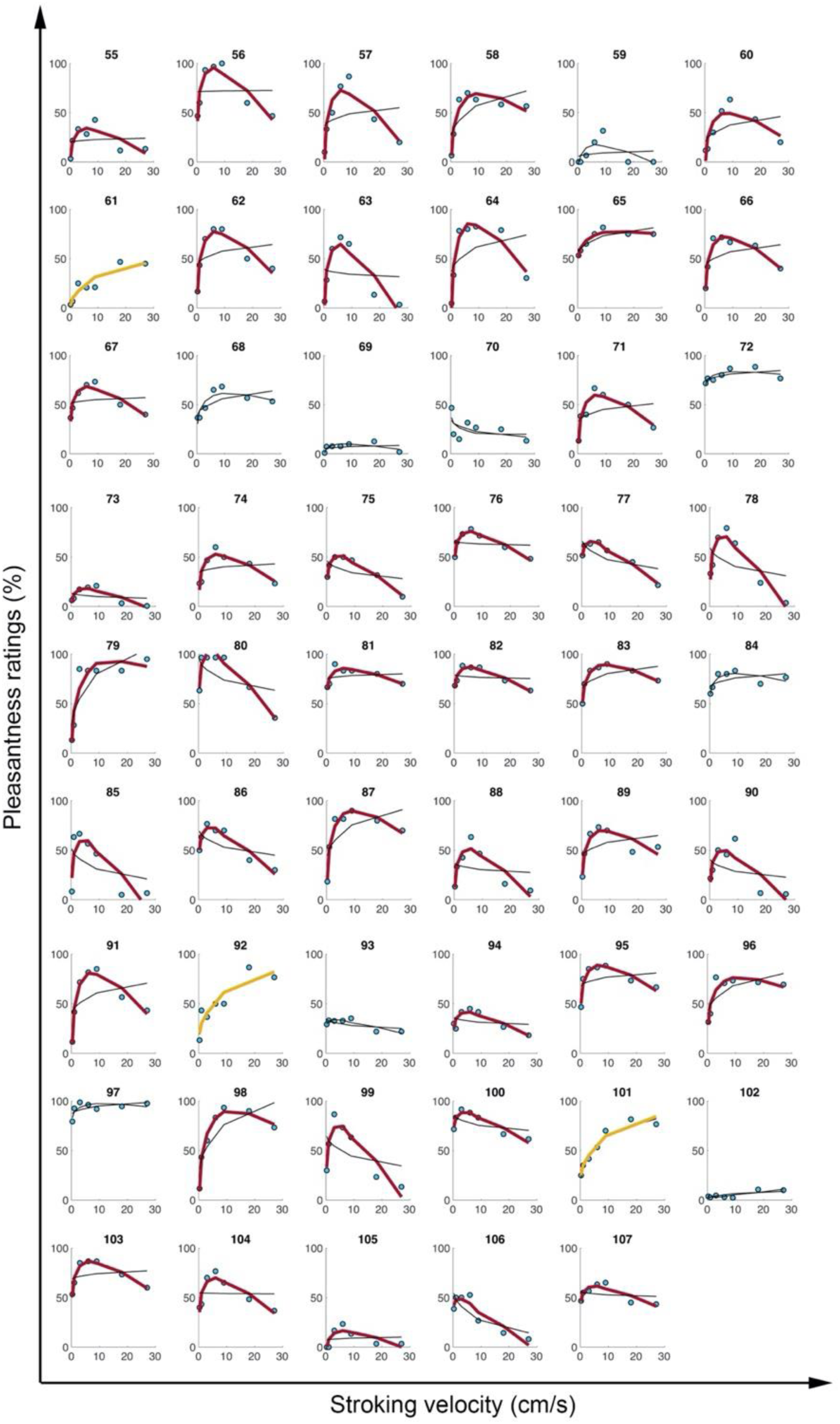
Individual pleasantness ratings in response to strokes applied to the forearm at different velocities (Experiment 1). The figure displays the individual pleasantness ratings at each stroking velocity (blue dots) and the linear and quadratic fit results (curves). A thick red curve indicates that the participant has a ’quadratic’ profile (i.e., significant quadratic fit and *p values* (LRT) < .05). A thick yellow curve indicates that the participant has a ‘linear’ profile (i.e., significant linear fit and *p values* (LRT) > .05). The absence of a bold curve (neither red nor yellow) indicates that the participant has a ‘random’ profile.

#### b. Sub-threshold psychopathology scales and cardiac interoceptive accuracy

First, we investigated whether the individual characteristics we collected could predict how well the quadratic model would predict the pleasantness ratings reported by the participants. We observed a small but significant negative correlation between the cardiac interoceptive accuracy score and the RMSE. For the whole population, the better the cardiac interoceptive accuracy was, the better the fit of the quadratic model (Table 2).

**Table 2:** Correlation results for the stroking delivered to the forearm.

| Variable of interest | Spearman correlation | S | P | Rho |
| --- | --- | --- | --- | --- |
| RMSE | EDE-Q | 39856 | 0.37 | 0.04 |
|  | Depression | 210404 | 0.62 | -0.03 |
|  | Anxiety | 202394 | 0.46 | 0.01 |
|  | **Cardiac interoceptive accuracy** | **236066** | **0.04** | **-0.16** |
| RSDd | EDE-Q | 39190 | 0.32 | 0.06 |
|  | Depression | 192923 | 0.29 | 0.06 |
|  | Anxiety | 205676 | 0.53 | -0.01 |
|  | Cardiac interoceptive accuracy | 212430 | 0.34 | -0.04 |

We then focused on the participants with a ‘quadratic’ profile. We investigated whether the individual characteristics correlated with the shape of the estimated curve, and more precisely, the A2 and A0 coefficients. We found that higher depression scores indicated a flatter curve: higher depression scores were associated with A2 coefficients closer to 0 (Table 3).

**Table 3:** Correlation results for the stroking delivered to the forearm for the participants with a ‘quadratic’ profile (N = 72).

| Variable of interest | Spearman correlation | S | P | Rho |
| --- | --- | --- | --- | --- |
| A2 | EDE-Q | 8206 | 0.27 | 0.1 |
|  | **Depression** | **49024** | **0.04** | **0.21** |
|  | Anxiety | 60932 | 0.43 | 0.02 |
|  | Cardiac interoceptive accuracy | 53088 | 0.89 | 0.15 |
| A0 | EDE-Q | 7406 | 0.87 | 0.19 |
|  | Depression | 55841 | 0.80 | 0.1 |
|  | Anxiety | 54604 | 0.85 | 0.12 |
|  | Cardiac interoceptive accuracy | 52114 | 0.09 | 0.16 |

### Pleasantness for different stroking velocities on the palm (non-hairy skin)

#### a. Categorization into ‘quadratic’, ‘linear’, or ‘random’ profiles

At the group level, we observed the classic inverted U-shaped curve. These classic replication results can be found in the Appendix 1 (see Appendix Figures 6 and 7). At the individual level, 28 participants (26.2%) did not show a significant effect of velocity on perceived pleasantness and were thus categorized as ‘random’. Out of the 79 remaining participants, 67 (62.6%) showed a ‘quadratic’ profile, and 11 (10.3%) showed a ‘linear’ profile (Figure 2). The raw sum for the RSD differences between the quadratic and linear models was -548, and the CI analysis confirmed that the quadratic regression outperformed the linear regression (lower bound: -642; upper bound: -457). Overall, our results showed once again that despite individual differences, the influence of stroking velocity on pleasantness ratings was better described by a quadratic regression than by a linear regression at the individual level as it was at the group level.

**Figure 2:**
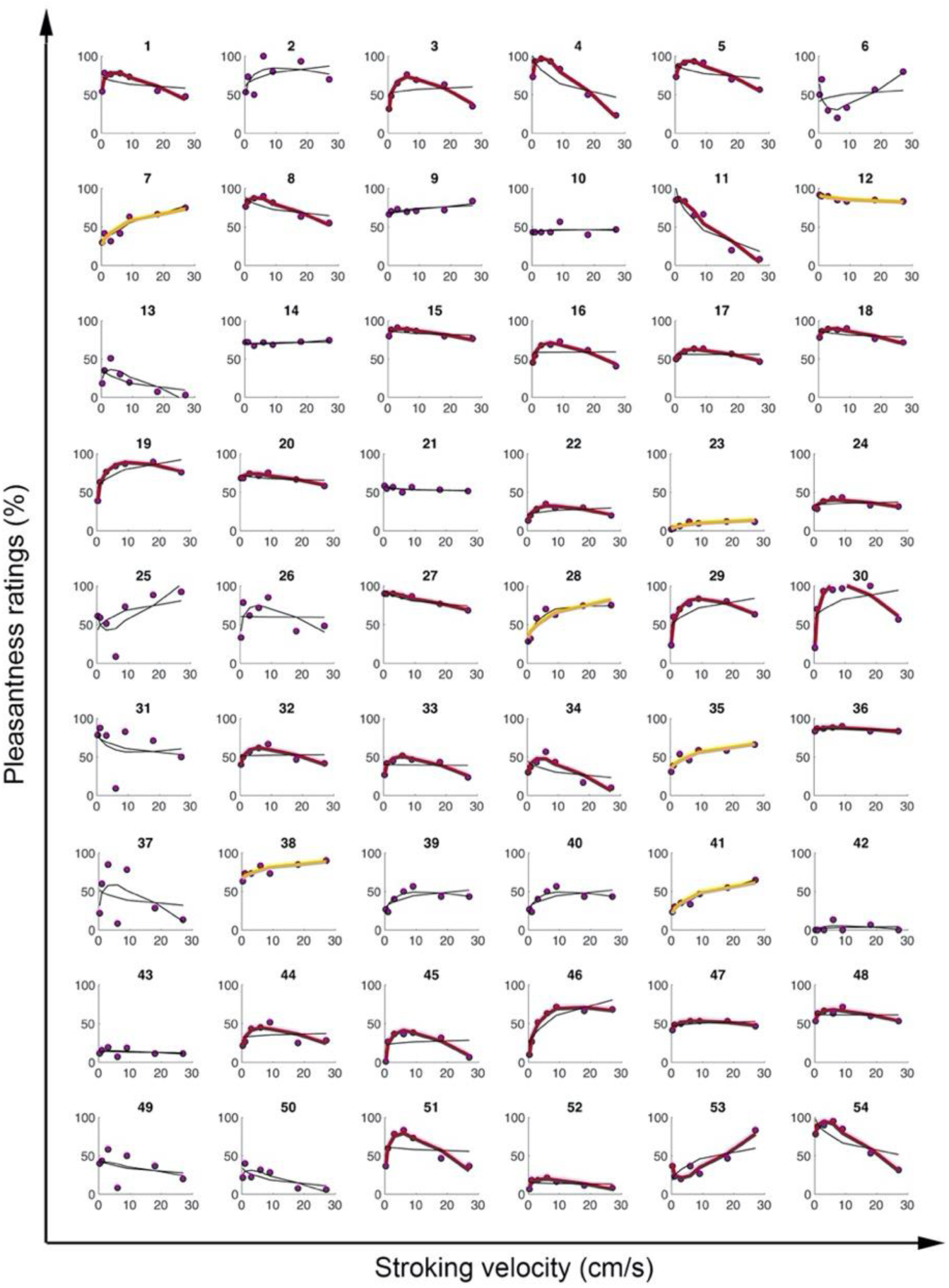
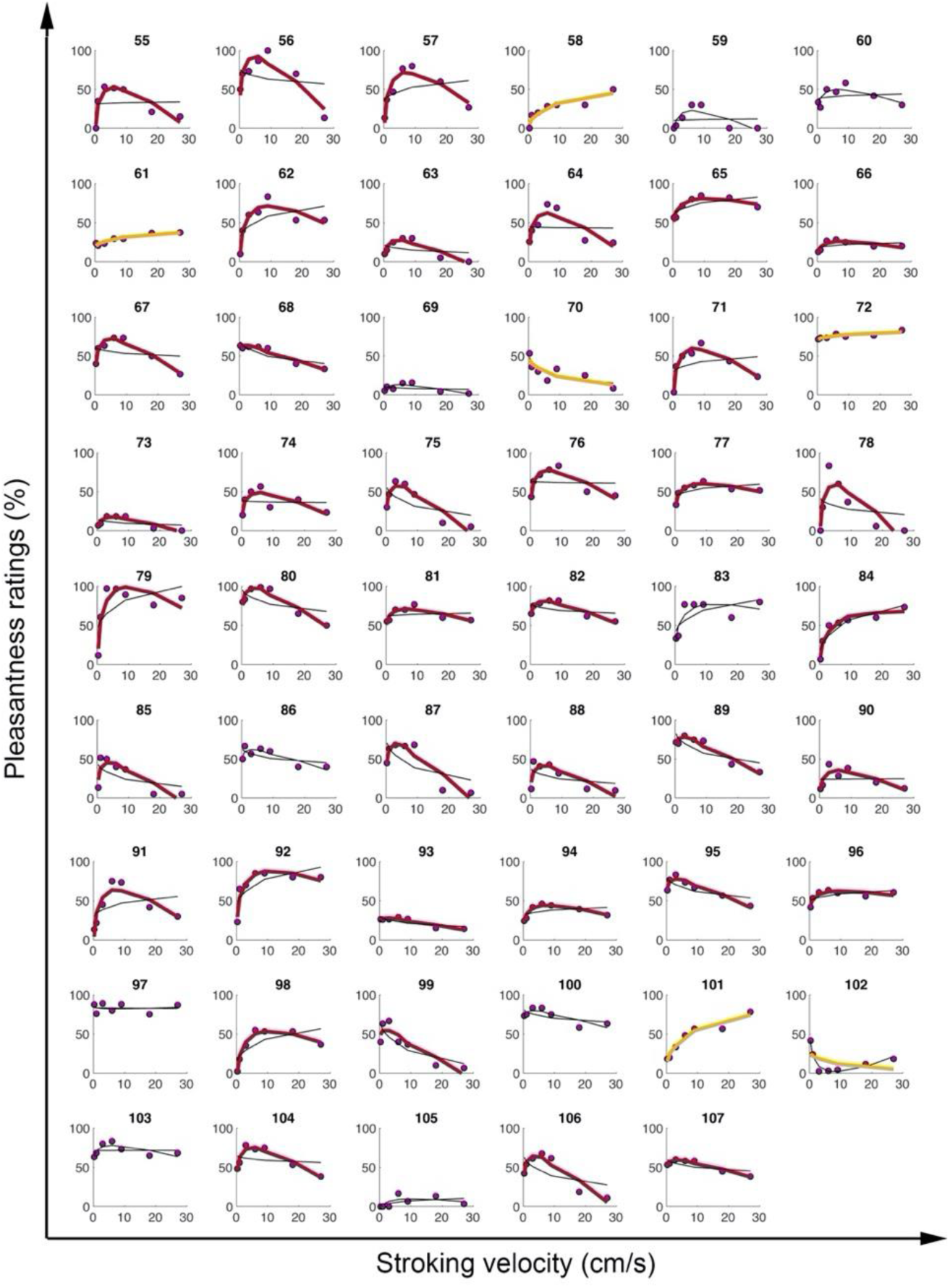
Quadratic and linear fit of individual pleasantness ratings in response to strokes applied to the palm at different velocities (Experiment 1). The figure displays the individual pleasantness ratings at each stroking velocity (purple dots) and the linear and quadratic fit results (curves). A thick red curve indicates that the participant has a ’quadratic’ profile (i.e., significant quadratic fit and *p values*(LRT) < .05). A thick yellow curve indicates that the participant has a ‘linear’ profile (i.e., significant linear fit and *p values*(LRT) > .05). The absence of a bold curve (neither red nor yellow) indicates that the participant has a ‘random’ profile.

#### b. Sub-threshold psychopathology scales and cardiac interoceptive accuracy

The following correlation analyses are the same as those reported before: we investigated whether the interoceptive indices we collected could predict the shape of the pleasantness ratings reported by the participants, but this time for stimulation delivered to the palm. For the whole population, no significant correlation was observed. Only trends were identified between interoceptive accuracy and RMSE (Table 4), while this correlation was significant for the data collected after stroking the forearm (Table 2).

**Table 4:**
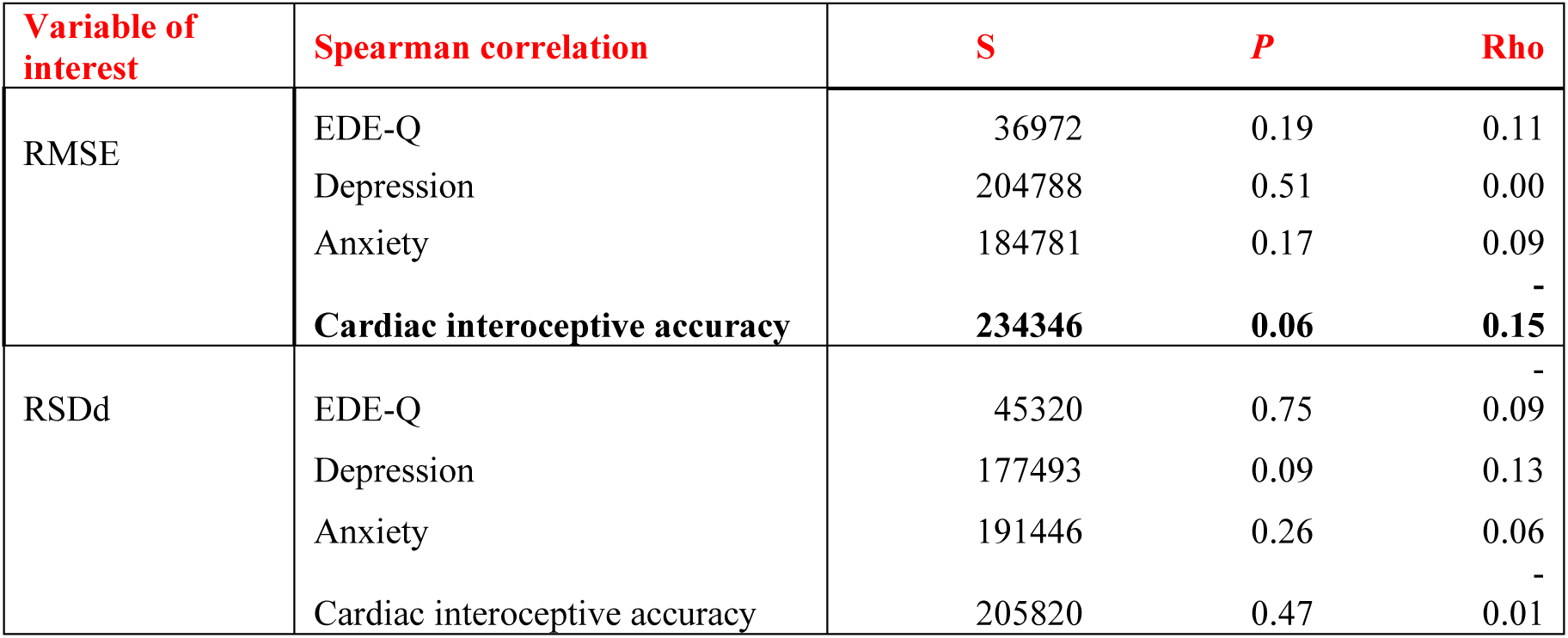
Correlation results for the stroking delivered to the palm.

We then performed the same analyses we performed for the forearm ratings, focusing only on the participants with a ‘quadratic’ profile and the A2/A0 coefficients. No significant correlations were observed (Table 5).

**Table 5:**
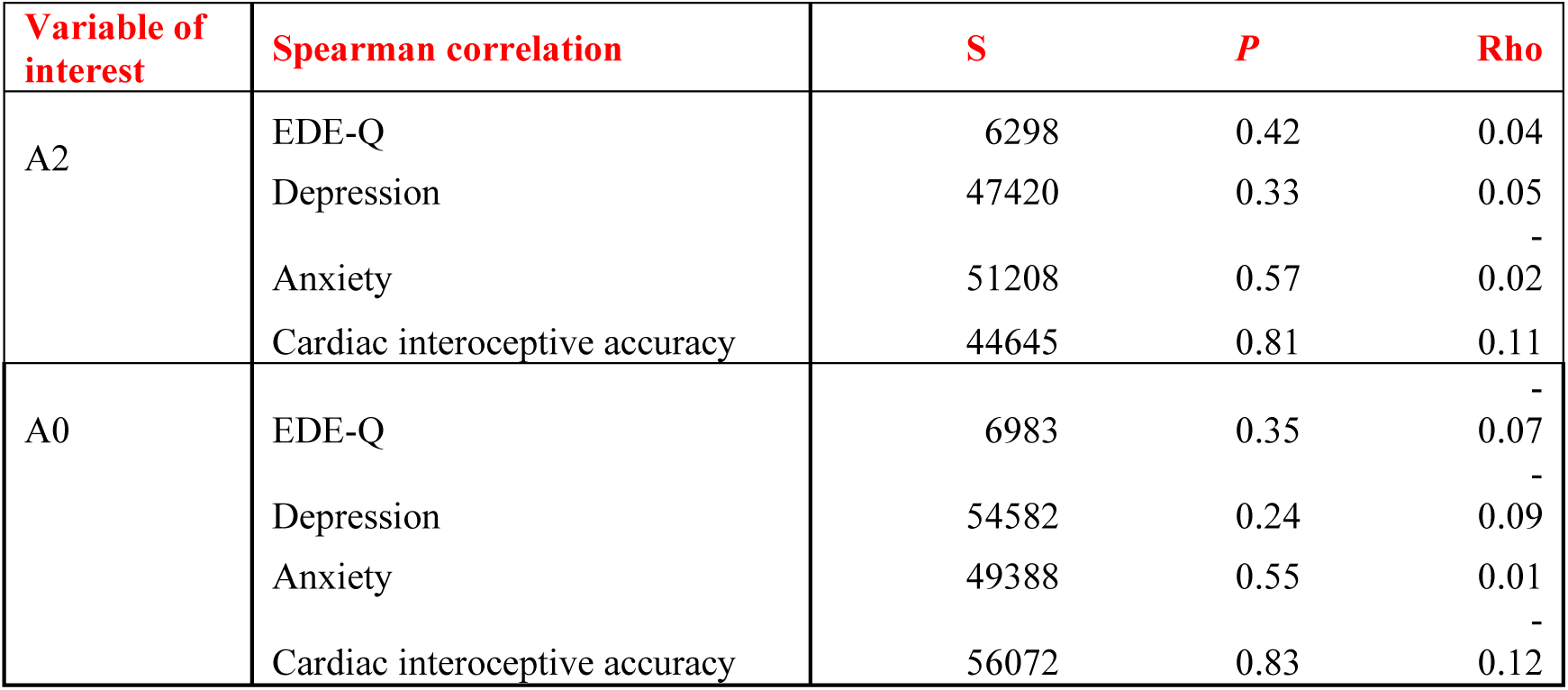
Correlation results for the stroking delivered to the forearm for the participants with a ‘quadratic’ profile (N = 67).

### Stability of pleasantness ratings across body sites

#### a. Repartition in profiles

Sixty-seven (62.6%) participants displayed the same profile for forearm and palm stroking. This result is above chance level (33%). The detailed repartition of the participants is shown in Table 6. Furthermore, the observed likelihood for each profile exceeded the predicted likelihood (Table 7). Again, this result suggested that the identified profiles were stable across body sites.

**Table 6:** Repartition of the participants into profiles for both body sites.

|  |  | Palm profiles |  |  |  |
| --- | --- | --- | --- | --- | --- |
|  |  | Quadratic | Linear | Random | Total |
| Forearm profiles | Quadratic | 53 | 4 | 15 | 72 |
|  | Linear | 3 | 4 | 3 | 10 |
|  | Random | 11 | 4 | 10 | 25 |
|  | Total | 67 | 12 | 28 | 107 |

**Table 7:** Predicted and observed likelihoods for each profile to be identical at both body sites.

|  | Predicted likelihood | Observed likelihood | Difference | Interpretation |
| --- | --- | --- | --- | --- |
| Quadratic | 0.453 | 0.495 | 0.043 | The observed likelihood is always higher than the predicted one. This is an argument in favor of stability. |
| Linear | 0.009 | 0.037 | 0.029 |  |
| Random | 0.055 | 0.093 | 0.039 |  |

#### b. Stability of the model’s coefficients for participants with a ‘quadratic’ profile

This section examines the correlations between the model coefficients A2 (quadratic term), A1 (linear term), and A0 (intercept) for the different body sites for the participants who had a ‘quadratic’ profile at both the forearm and the palm. Each coefficient was significantly correlated with its homolog from the other body site (A2: S = 8571, *p* < .001, rho = 0.65; A1: S = 8604, *p* < .001, rho = 0.65; A0: S = 4970, *p* < .001, rho = 0.80; see Appendix 1, Appendix Figure 7). These strong observed correlations argue in favor of a stability in pleasantness judgment across body sites. Too few participants (four) showed a stable “linear” profile for a similar analysis in this subpopulation to be meaningful.

## EXPERIMENT 2

The focus of Experiment 2 was to test the stability of the perceived pleasantness across two experimental sessions performed a week apart. We compared the observed and predicted likelihood for a participant to be in the same profile category in both sessions. The predicted likelihood was calculated as the square of the probability of having a given profile in session 1. The observed likelihood was calculated as the product of the probability of having a given profile in session 1 multiplied by the probability of participants with this given profile also having this profile in session 2. This analysis is similar to the one used in Experiment 1 to assess the stability across body sites and by Bendas et al. (2021) to assess the stability of pleasantness ratings across repetitions on two different days. Again, the participants who had a ‘quadratic’ profile in both sessions could still display very different patterns of response (e.g., steep versus flat curves, different maximum and minimum). Thus, we investigated the correlation between the coefficients of the quadratic model, A2, A1, and A0, from one session to the other.

### Method

#### Participants and procedure

Forty-one participants from Experiment 1 took part in this follow-up (20 women and 21 men; 25.1 ± 4 years). These participants underwent the same exact procedure as in Experiment 1 twice in total, with the two sessions being one week apart. This group was also tested at another hairy-skin body site, i.e., the back of their hand (dorsal condition), in both sessions. This body site is less often investigated than the forearm; therefore, we chose to report mainly the comparison between the palm and forearm. However, the results regarding the dorsal condition can be found in the Appendix 2 (Appendix Figure 13).

#### Data analysis

For each body site, the data from the two sessions were analyzed separately. The model fitting and profile categorization were the same as those used in Experiment 1.

### Results

#### Response to forearm (hairy skin) stimulation

##### a. Quadratic, linear, and random profile categorization

At the group level, we observed the classic inverted U-shaped curve. These results replicating the classically observed group-level results for tactile pleasantness ratings can be found in the Appendix 2 (see Appendix Figure 9). In response to stimulation applied to the forearm, 8 (19.5%) participants did not show a significant effect of velocity on perceived pleasantness and were thus categorized as ‘random’, 31 (75.6%) participants were categorized as having a ‘quadratic’ profile, and 2 (4.9%) participants were categorized as having a ‘linear’ profile in session 1 (Figure 3). In session 2, 7 (17.1%) participants were categorized as having a ‘random’ profile, 28 (68.3%) participants were categorized as having a ‘quadratic’ profile, and 6 (14.6%) participants were categorized as having a ‘linear’ (Figure 3). As in Experiment 1, the CI analysis showed a clear superiority of the quadratic model for both sessions (session 1: lower bound: -356, raw sum: -291, upper bound: -226; session 2: lower bound: - 310, raw sum: -255, upper bound: -202).

**Figure 3:**
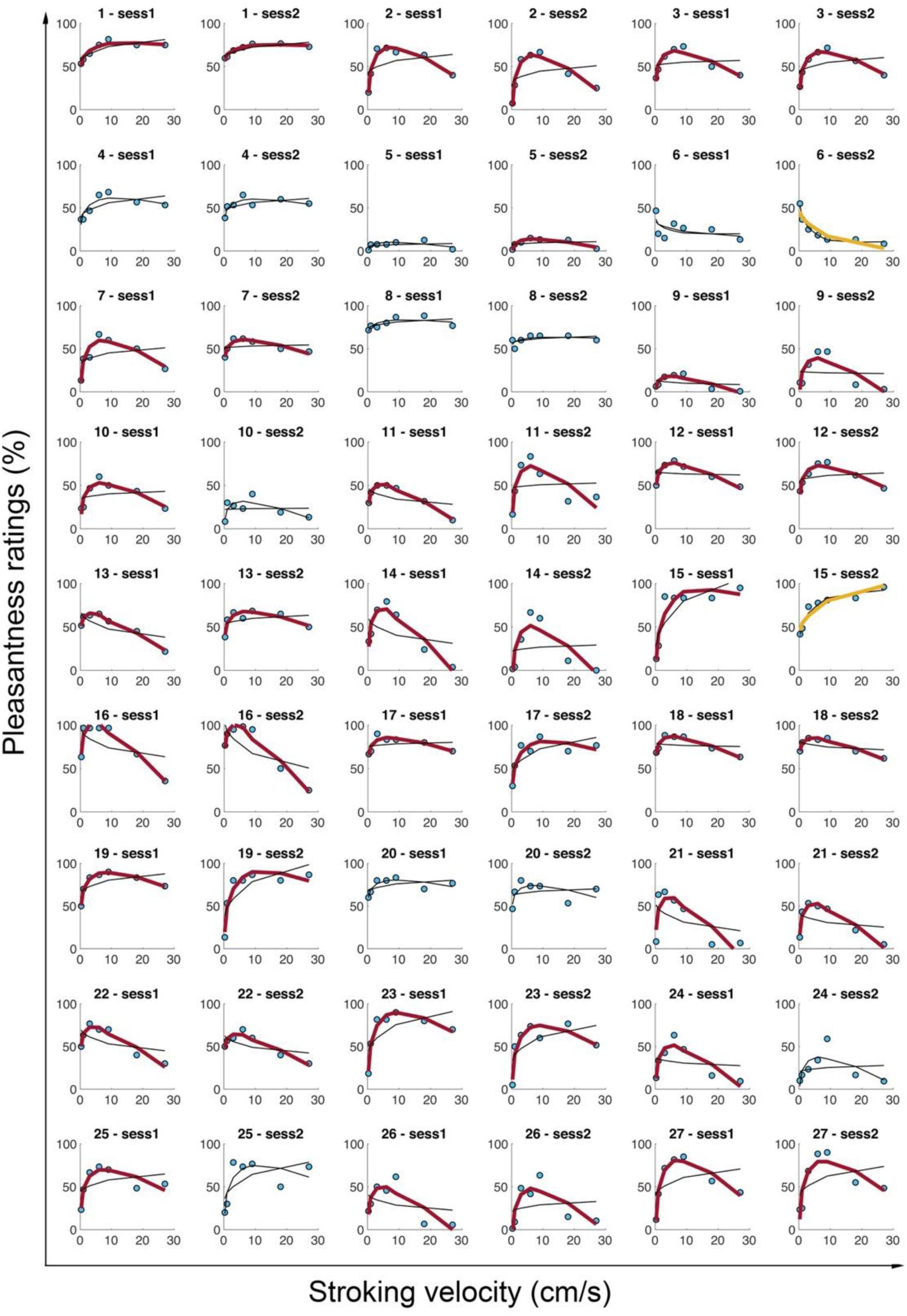
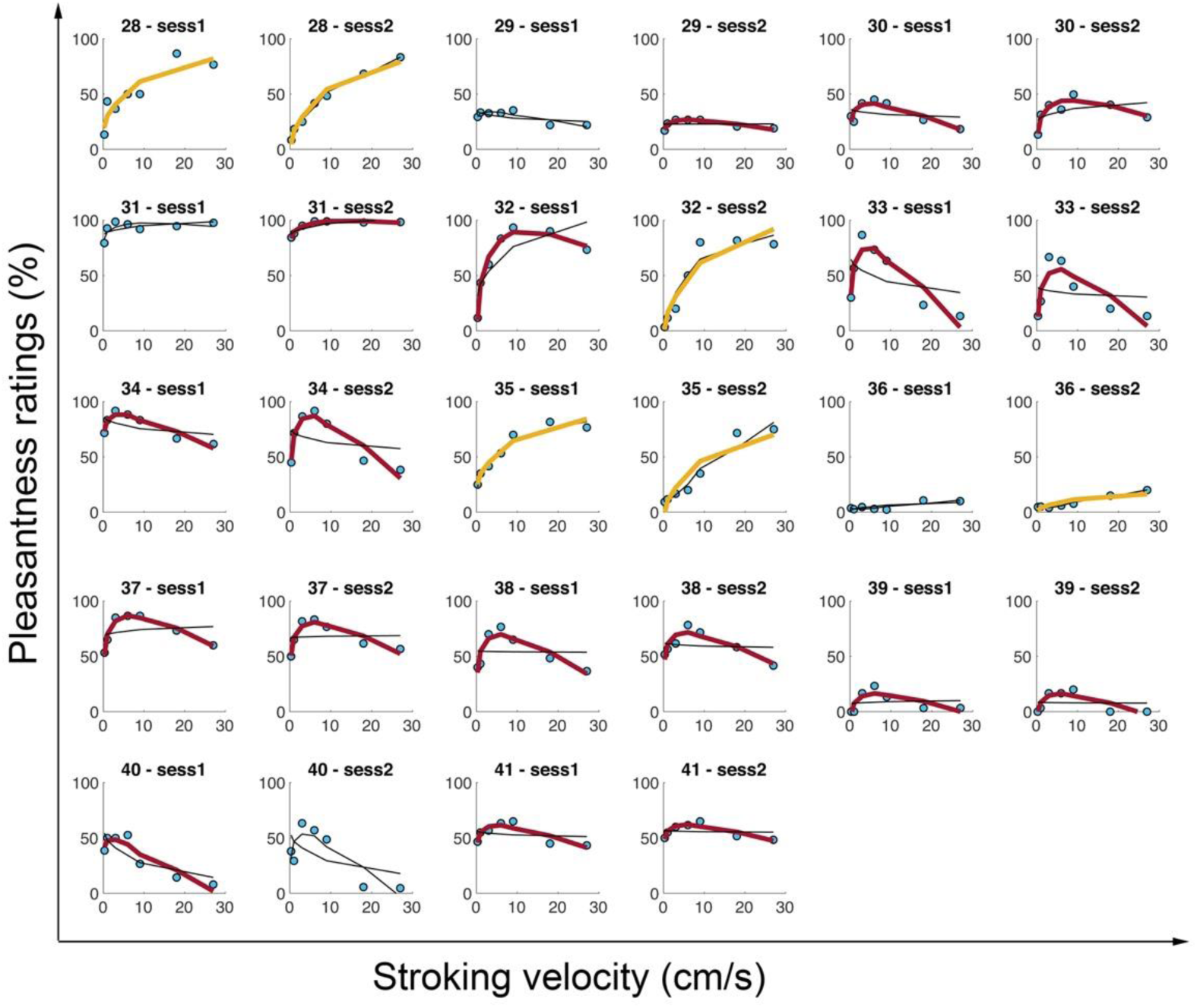
Quadratic and linear fit of individual pleasantness ratings in response to strokes applied to the forearm for both sessions (Experiment 2). The figure displays the individual pleasantness ratings at each stroking velocity (blue dots) and the linear and quadratic fit results (curves). A thick red curve indicates that the participant has a ’quadratic’ profile (i.e., significant quadratic fit and *p values*(LRT) < .05). A thick yellow curve indicates that the participant has a ‘linear’ profile (i.e., significant linear fit and *p values*(LRT) > .05). The absence of a bold curve (neither red nor yellow) indicates that the participant has a ‘random’ profile. For each participant, the results from the two sessions are plotted side-by-side.

##### b. Stability across sessions for forearm stroking

Twenty-five (73%) participants displayed the same profile in session 1 and session 2. This result was above chance level (33%). The detailed repartition of the participants is shown in Table 8. Furthermore, the observed likelihood for each profile exceeded the predicted likelihood (Table 9). This result suggested that the identified profiles were stable across sessions.

**Table 8:** Repartition of the participants into profiles for both sessions.

|  |  | Session 2 profiles |  |  |  |
| --- | --- | --- | --- | --- | --- |
|  |  | Quadratic | Linear | Random | Total |
| Session 1 profiles | Quadratic | 25 | 2 | 4 | 31 |
|  | Linear | 0 | 2 | 0 | 2 |
|  | Random | 3 | 2 | 3 | 8 |
|  | Total | 28 | 6 | 7 | 41 |

**Table 9:** Predicted and observed likelihoods for each profile to be identical in both sessions.

|  | Predicted likelihood | Observed likelihood | Difference | Interpretation |
| --- | --- | --- | --- | --- |
| Quadratic | 0.57 | 0.61 | 0.04 | The observed likelihood is higher than the predicted one. This is an argument in favor of stability |
| Linear | 0.00 | 0.05 | 0.05 |  |
| Random | 0.04 | 0.07 | 0.04 |  |

We then examined the correlations between the model coefficients A2 (quadratic term), A1 (linear term), and A0 (intercept) between the two sessions for the 25 participants who had a ‘quadratic’ profile in both sessions. Each coefficient was significantly correlated with its homolog from the other session (A2: S = 886, *p* < .001, rho = 0.66; A1: S = 817, *p* < .001, rho = 0.69; A0: S = 802, *p* < .001, rho = 0.69; see Appendix 2, Appendix Figure 10). These strong observed correlations argue in favor of a stability of pleasantness judgment across sessions.

#### Response to palm (non-hairy skin) stimulation

##### a. Quadratic, linear, and random profile categorization

At the group level, we observed the classic inverted U-shaped curve. These results replicating the classically observed group-level results for tactile pleasantness ratings can be found in the Appendix 2 (see Appendix Figure 11). In response to stimulation applied to the palm, 8 (19.5%) participants did not show a significant effect of velocity on perceived pleasantness and were thus categorized as having a ‘random’ profile, 30 (73.1%) participants were categorized as having a ‘quadratic’ profile, and 3 (7.3%) participants were categorized as having a ‘linear’ profile in session 1 (Figure 4). In session 2, 7 (17.1%) participants were categorized as having a ‘random’ profile, 28 (68.3%) participants were categorized as having a ‘quadratic’ profile, and 6 (14.6%) participants were categorized as having a ‘linear’ profile (Figure 4). As in Experiment 1, the CI analysis showed a clear superiority of the quadratic model for both sessions (session 1: lower bound: - 345, raw sum: - 285, upper bound: -226; session 2: lower bound: - 336, raw sum: - 276, upper bound: -219).

**Figure 4:**
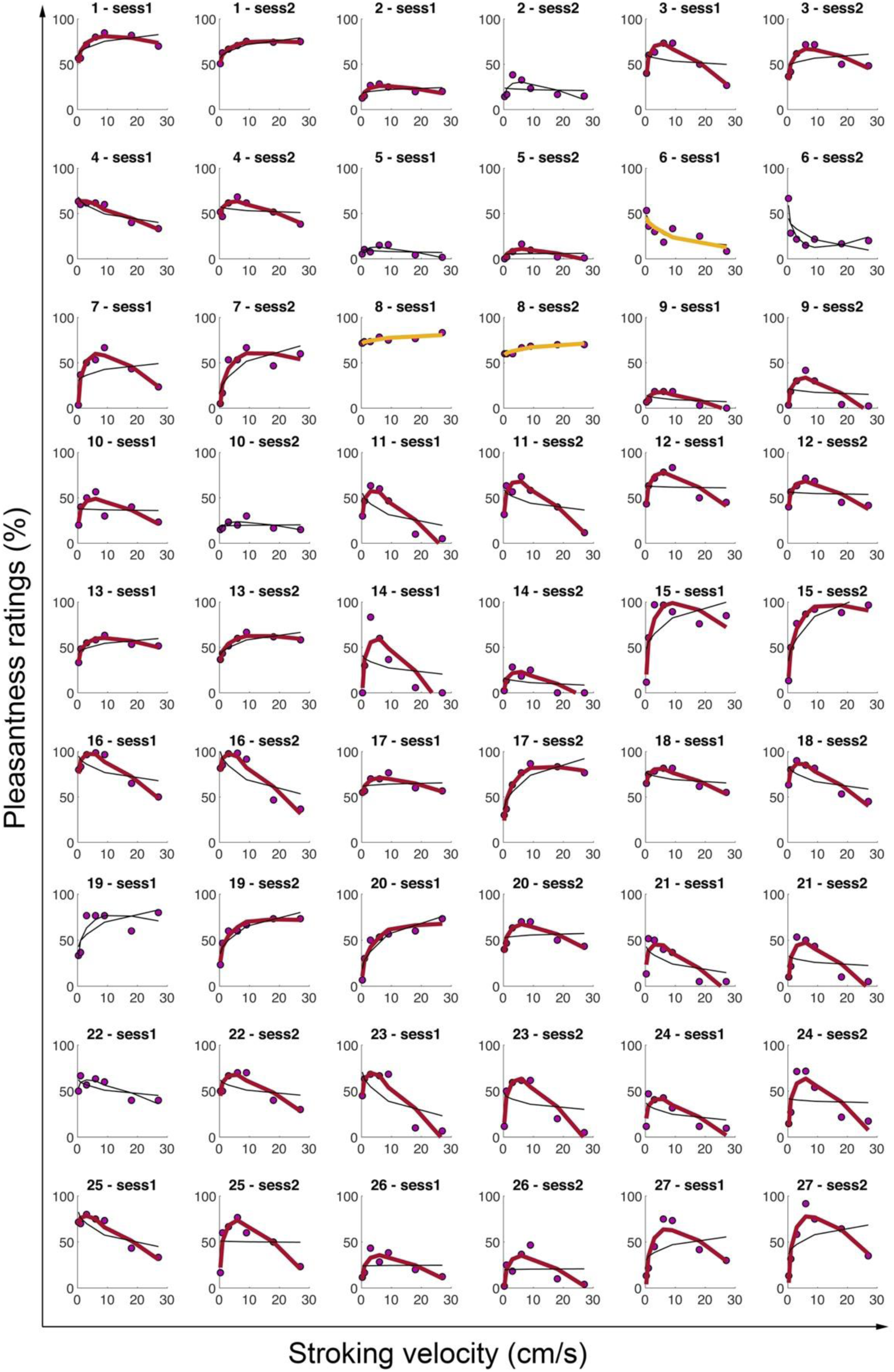
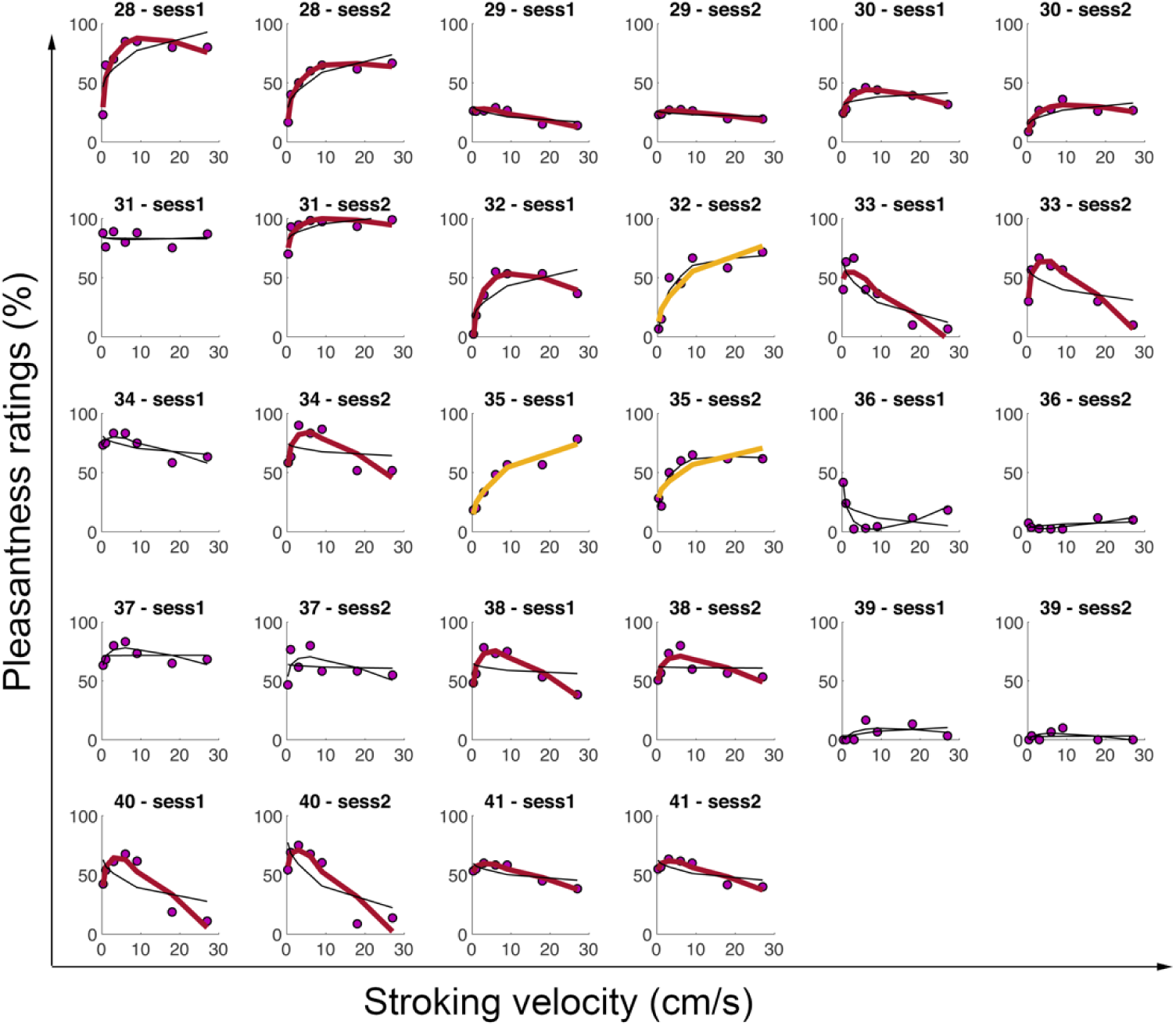
Quadratic and linear fit of individual pleasantness ratings in response to strokes applied to the palm for both sessions (Experiment 2). The figure displays the individual pleasantness ratings at each stroking velocity (purple dots) and the linear and quadratic fit results (curves). A thick red curve indicates that the participant has a ’quadratic’ profile (i.e., significant quadratic fit and *p values*(LRT) < .05). A thick yellow curve indicates that the participant has a ‘linear’ profile (i.e., significant linear fit and *p values*(LRT) > .05). The absence of a bold curve (neither red nor yellow) indicates that the participant has a ‘random’ profile. For each participant, the results from the two sessions are plotted side-by-side.

##### b. Stability across sessions for palm stroking

Thirty-two (78%) participants displayed the same profile in session 1 and session 2. This result was above chance level (33%). The detailed repartition of the participants is shown in Table 10. Furthermore, the observed likelihood for each profile exceeded the predicted likelihood (Table 11). This result suggested that the identified profiles were stable across sessions.

**Table 10:** Repartition of the participants into profiles for both sessions.

|  |  | Session 2 profiles |  |  |  |
| --- | --- | --- | --- | --- | --- |
|  |  | Quadratic | Linear | None | Total |
| Session 1 profiles | Quadratic | 27 | 1 | 2 | 30 |
|  | Linear | 0 | 2 | 1 | 3 |
|  | Random | 5 | 0 | 3 | 8 |
|  | Total | 32 | 3 | 6 | 41 |

**Table 11:** Predicted and observed likelihoods for each profile to be identical in both sessions.

|  | Predicted likelihood | Observed likelihood | Difference | Interpretation |
| --- | --- | --- | --- | --- |
| Quadratic | 0.54 | 0.66 | 0.12 | The observed likelihood is always higher than the predicted one. This is an argument in favor of stability |
| Linear | 0.01 | 0.05 | 0.04 |  |
| Random | 0.04 | 0.07 | 0.04 |  |

We then examined the correlations between the model coefficients A2 (quadratic term), A1 (linear term), and A0 (intercept) between the two sessions for the 27 participants who had a ‘quadratic’ profile in both sessions. Each coefficient was significantly correlated with its homolog from the other session (A2: S = 1668, *p* < .05, rho = 0.49; A1: S = 2010, *p* < .05, rho = 0.39; A0: S = 1139, *p* < .001, rho = 0.65; see Appendix 2, Appendix Figure 12). These observed correlations argue in favor of a stability of pleasantness judgment across sessions. However, the correlation between A1 in session 1 and session 2 was weaker than the correlations previously reported for forearm stimulation.

## EXPERIMENT 3

The focus of Experiment 3 was to test the stability of the perceived pleasantness across several repetitions. Sensory processing is “noisy”, and this intrinsic uncertainty in our sensory system may explain differences between individuals and differences between studies (e.g., Croy, Bierling, Sailer & Ackerley, 2021). In Experiments 1 and 2, we found a higher percentage of individuals for whom the quadratic regression was significant than in a previous study (67.1% vs. 42%; Croy et al., 2021). Most of the studies assessing pleasantness in response to different velocities of stroking assess each condition 3 times. This low number of repetitions is justified by the quick habituation and fatigue of affective touch fibers. Triscoli, Ackerley & Sailer (2014) showed that tactile pleasantness diminished during 50 minutes of repetitions, particularly when touch was delivered at 3 cm/s, compared to slower (0.3 cm/s) and faster (30 cm/s) velocities. Nevertheless, when interested in individual perception, an optimal approach would be to collect more data per participant to better account for individual variability and sensory uncertainty (MacDonald, Nyberg, & Bäckman, 2006). In Experiment 3, we propose a new protocol that includes more repetitions in each condition to better account for individual variability. By doing so, our aim was to evaluate whether the number of repetitions influenced the categorization of participants into profiles (quadratic, linear, or random). We first compared the profile repartition we obtained with the classic three repetitions of each stroking velocity (as we did in the first two experiments) to the results obtained with ten repetitions. Then, for a more fine-grained evaluation of the impact of the number of repetitions on the shape of the tactile pleasantness ratings, we focused on how the fit of the quadratic model and the corresponding estimated parameters varied with increasing repetitions.

### Method

#### Participants

Sixteen healthy, naïve participants took part in Experiment 3 (6 women; 24.3 ± 5 years). All volunteers provided written informed consent prior to their participation. All participants received a cinema ticket as compensation for each hour spent on the experiment. All experiments were approved by the Swedish Ethics Review Authority.

#### Affective touch task procedure

We followed the same procedure as in Experiment 1. However, each velocity was presented ten times, for a total of 70 stroking trials per location (palm and forearm, in randomized order).

#### Data analysis

##### a. Descriptive modeling: Quadratic, linear, and random profile categorization

The model fitting and profile categorization were the same as in Experiments 1 and 2. We compared the outcomes when the fit was performed on all data or when considering the first 3 repetitions in each condition.

##### b. Influence of the number of repetitions

As shown in the previous two experiments, the negative quadratic model seemed to be an efficient model to capture the perception of pleasantness in response to touch delivered at different velocities at the individual level. However, are the fitting outcomes significantly impacted by the number of repetitions of each type of tactile stimulation? To answer this question, we fitted the quadratic model to the participants’ pleasantness ratings for the first presentation of each velocity only and then repeated the same fitting procedure taking into account one additional presentation of each velocity each time (from 2 to 10 repetitions). Thus, we obtained 10 fitting outcomes per participant. We used linear mixed models to evaluate the impact of the number of repetitions at each stroking velocity on the goodness of fit of the quadratic model (reflected by the RMSE) and on the shape of the fitting curve (reflected by the estimated A2 and A0 coefficients).

#### Results

##### a. Quadratic, linear, and random profile categorization

At the group level, we observed the classic inverted U-shaped curve (see Appendix 2, Appendix Figure 16). In this experiment, similar to Experiments 1 and 2, the ‘quadratic’ profile was predominant. Based on Triscoli, Ackerly and Sailer (2014), we expected that this predominance might be reduced when considering the 10 repetitions for each condition: with more repetitions, the sensory attenuation would probably increase, and the participants might become less sensitive and more “numb” to the pleasantness, which could decrease the relevance of the quadratic model. However, the results actually suggested the opposite: regardless of whether stroking was applied to the forearm or the palm, a clear majority of participants presented a ‘quadratic’ profile after three repetitions at each velocity (forearm = 56%; palm: 75%); this relatively higher prevalence of the ‘quadratic’ profile further increased when each stimulation was repeated 10 times (forearm = 75%; palm: 94%), as the observed likelihood for this profile was higher than the predicted likelihood (Tables 12 and 13 and Figure 5). This suggested that the quadratic velocity-pleasantness response function was robust over repeated stimulations and that pleasant tactile sensations did not quickly attenuate.

**Figure 5:**
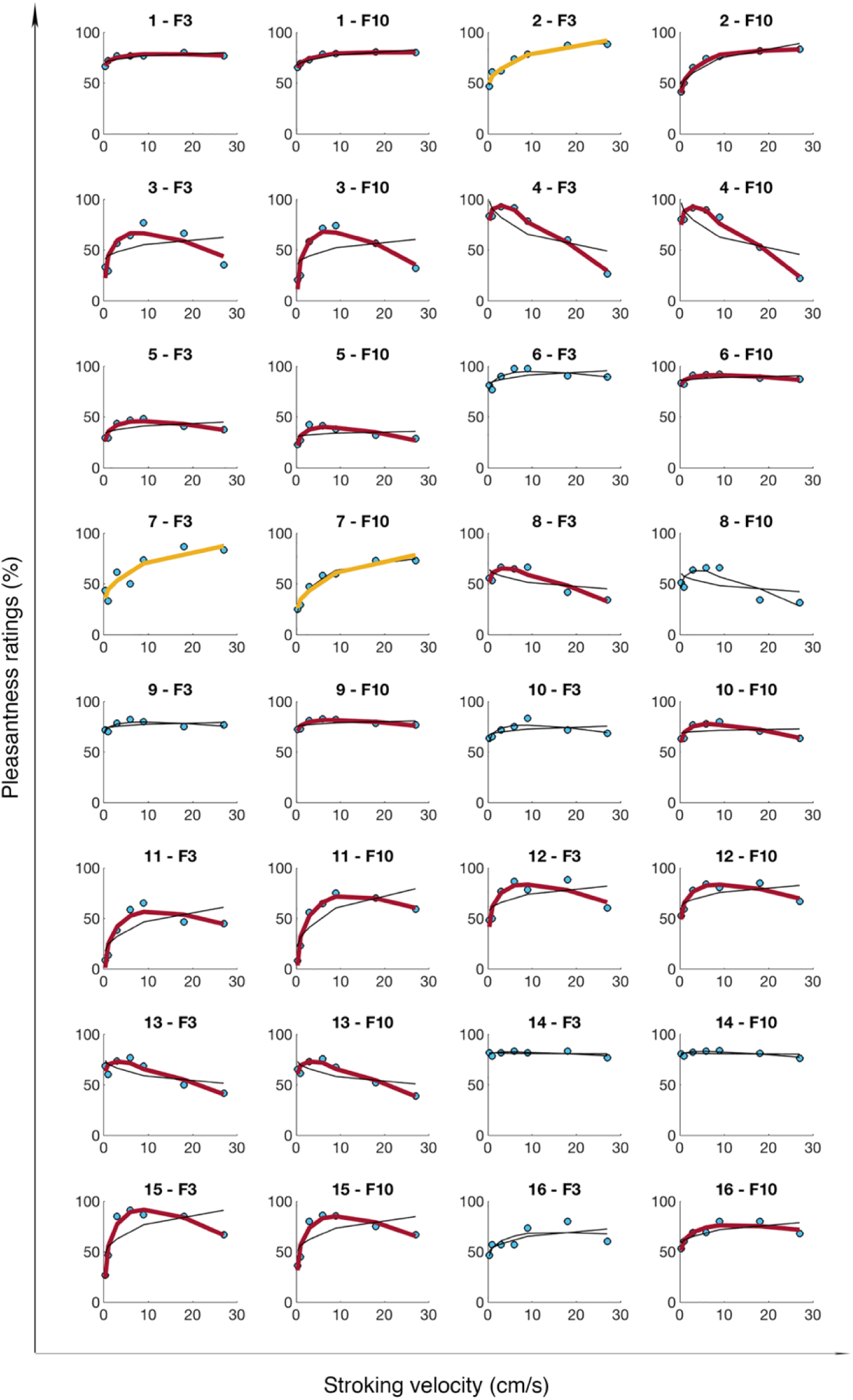
Quadratic and linear fit of individual pleasantness ratings in response to strokes applied to the forearm for 10 and 3 repetitions in each condition (Experiment 3). The figure displays the individual pleasantness ratings at each stroking velocity (blue dots) and the linear and quadratic fit results (curves). A thick red curve indicates that the participant has a ’quadratic’ profile (i.e., significant quadratic fit and *p-values*(LRT) < .05). A thick yellow curve indicates that the participant has a ‘linear’ profile (i.e., significant linear fit and *p values*(LRT) > .05). The absence of a bold curve (neither red nor yellow) indicates that the participant has a ‘random’ profile. For each participant, the results for 3 repetitions are plotted first, and the results when 10 repetitions are taken into account are plotted side-by-side.

**Figure 6:**
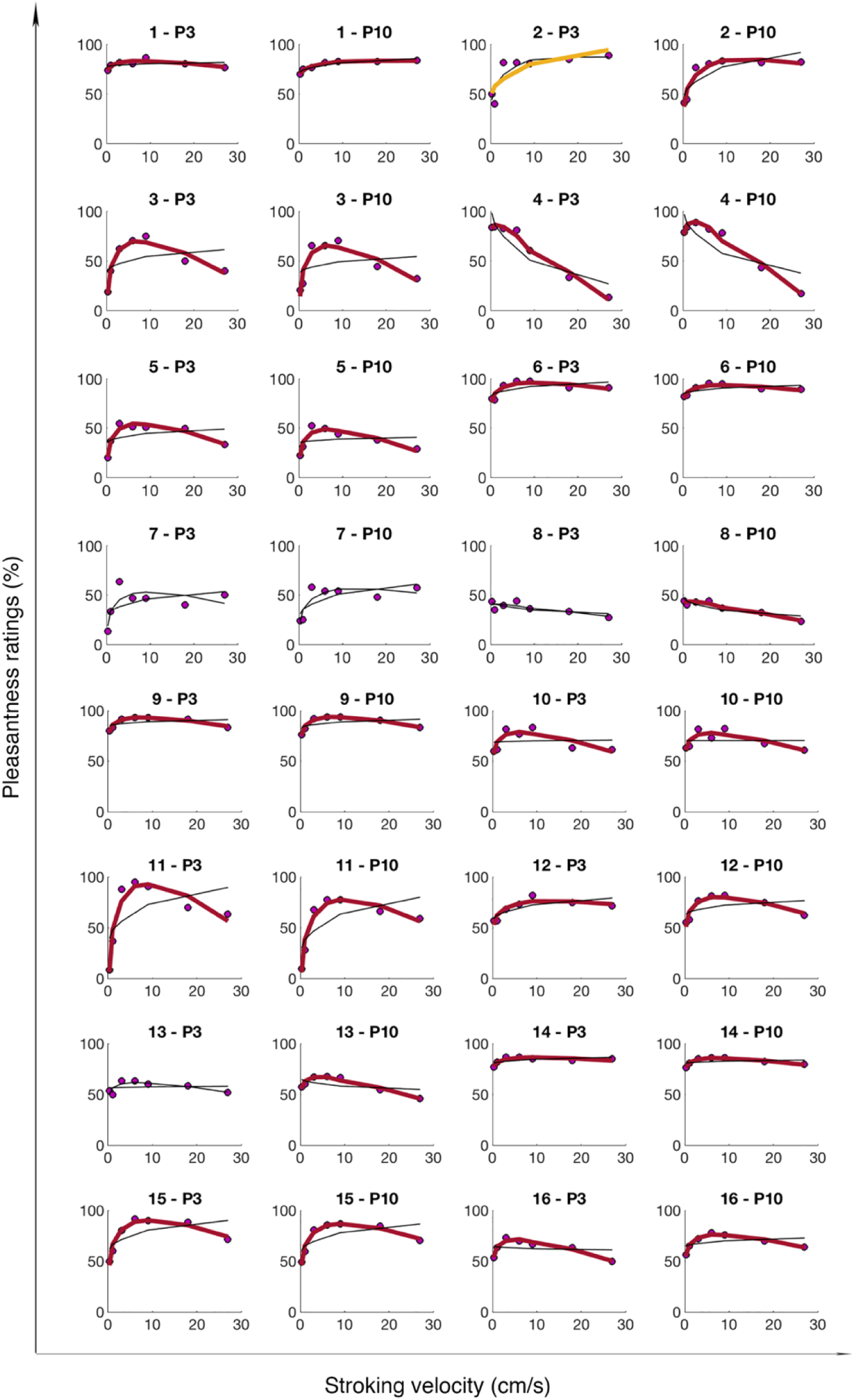
Quadratic and linear fit of individual pleasantness ratings in response to strokes applied to the palm for 10 and 3 repetitions in each condition (Experiment 3). The figure displays the individual pleasantness ratings at each stroking velocity (purple dots) and the linear and quadratic fit results (curves). A thick red curve indicates that the participant has a ’quadratic’ profile (i.e., significant quadratic fit and *p values*(LRT) < .05). A thick yellow curve indicates that the participant has a ‘linear’ profile (i.e., significant linear fit and *p values*(LRT) > .05). The absence of a bold curve (neither red nor yellow) indicates that the participant has a ‘random’ profile. For each participant, the results for 10 repetitions are plotted first, and the results when only 3 repetitions are taken into account are plotted side-by-side.

**Table 12:** Repartition of the participants into profiles when considering 10 or 3 repetitions in each condition.

| Forearm profiles |  |  |  |  |  | Palm profiles |  |  |  |  |  |
| --- | --- | --- | --- | --- | --- | --- | --- | --- | --- | --- | --- |
|  |  | 3 repetitions |  |  |  |  |  | 3 repetitions |  |  |  |
|  |  | Quadratic | Linear | None | Total |  |  | Quadratic | Linear | None | Total |
| 10 rep. | Quadratic | 8 | 0 | 4 | 12 | 10 rep. | Quadratic | 12 | 1 | 2 | 15 |
|  | Linear | 0 | 1 | 0 | 1 |  | Linear | 0 | 0 | 0 | 0 |
|  | Random | 1 | 1 | 1 | 3 |  | Random | 0 | 0 | 1 | 1 |
|  | Total | 9 | 2 | 5 | 16 |  | Total | 12 | 1 | 3 | 16 |

**Table 13:**
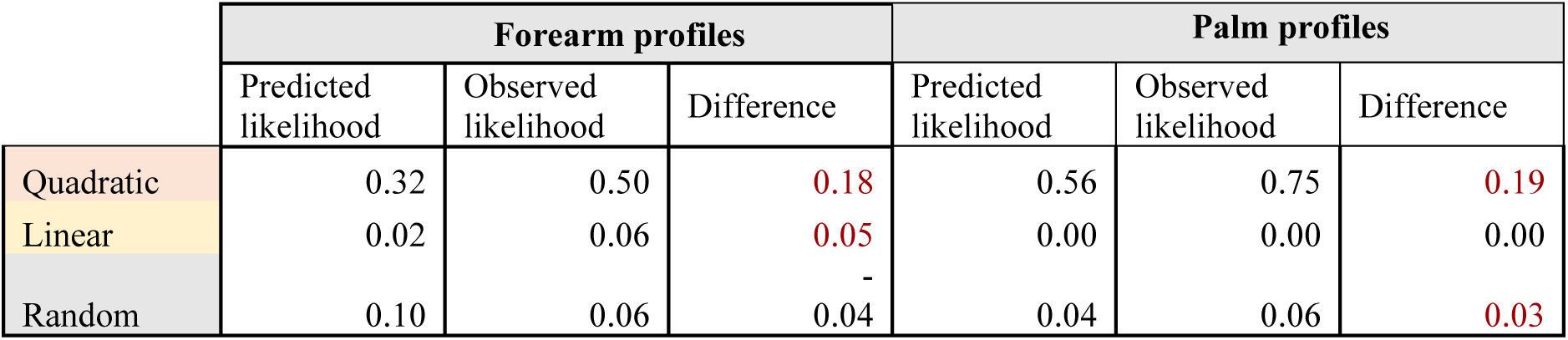
Predicted and observed likelihoods for each profile to be identical when considering the first 3 and then 10 repetitions at each stroking velocity.

##### b. Influence of the number of repetitions on fitting outcomes

By increasing the number of repetitions for each stroking velocity delivered to the forearm, we observed a significant decrease in RMSE values, i.e., the fit of the quadratic model became better with more repetitions (*F*(1,15)=26.8, *p* < .001), although the magnitude of this improvement remained modest (linear coefficient for the fixed effect of the number of repetitions: -0.31). However, the corresponding estimated A2 reflecting the curvature of the quadratic regression did not significantly change when the number of repetitions increased (*F*(1,15)=0.9, *p* = .37). A0, the coefficient representing the overall pleasantness perception did not seem to be significantly influenced by the number of repetitions (*F*(1,15)=1.5, *p* = .24; see Appendix 3, Appendix Figure 17).

By increasing the number of repetitions for each stroking velocity delivered to the palm, we observed no significant change in RMSE values, i.e., the fit of the quadratic model remained similar despite more repetitions (*F(*1,15)=0.4, *p* = .52). Once again, the corresponding estimated A2 coefficient reflecting the curvature of the quadratic regression did not significantly change when the number of repetitions increased (*F*(1,15)=1.5, *p* = .24). A0, the coefficient representing the overall pleasantness perception seemed to slightly increase with the number of repetitions (*F*(1,15)=5.6, *p* = .03; linear coefficient for the fixed effect of the number of repetitions: 0.89; see Appendix 3, Appendix Figure 18).

## Discussion

### Summary of key findings

Across three experiments, we 1) tested the hypothesis that the classic negative quadratic model (inverted-U shape function) would outperformed a linear model in describing the relationship between velocity of touch and subjective tactile pleasantness at the individual level in the majority of individuals; 2) explored how individual difference factors predicted the outcome of such individual fit (Experiment 1); 3) tested whether the shape of the quadratic model would be more pronounced and observed in more people on hairy compared to non-hairy skin, and 4) establish whether the individual shape of the tactile pleasantness-velocity functions was temporally stable, both across two identical experimental sessions one week apart (Experiment 2) and when the number of repetitions at each velocity increased (Experiment 3).

Our results showed that at the group level (N = 123), the relationship between the velocity of touch and subjective pleasantness was better described by a negative quadratic model than a linear model in the majority of participants. At the individual level, most participants (67.1% for the hairy skin site and 62.6% for the non-hairy skin site) showed a significant negative quadratic relationship between velocity of touch and pleasantness, which is substantially more frequent than previously reported (Croy, Bierling, Sailer & Ackerley, 2021) and thus more in line with the CT-hypothesis and suggesting that the inverted-U-shape function best captures individuals’ typical perception of affective touch. Unexpectedly, the frequency of the negative quadratic model fit, and the shape of this model, was similar across hairy (forearm) and non-hairy skin (palm) with 62.6% of participants displayed the same pattern at the forearm and palm. This result does not fit with the CT-hypothesis, but is more in line with more recent proposals that suggest that affective touch sensations arise from multiple afferent and top-down sources rather than being driven exclusively (or mainly) by afferent CT information. In terms of individual differences, we found that higher cardiac interoceptive accuracy was related to a better fit of the quadratic model of the perception of touch on hairy skin (and with a statistical trend for such relationship on non-hairy skin), and individual differences in self-reported depression influenced the steepness of the curve. Noteworthy, the patterns of the relationship between the velocity of touch and subjective pleasantness were temporally stable across two experimental sessions (73% of participants displayed the same profile across sessions on hairy skin and 78% on non-hairy skin; Experiment 2) and were robust and replicable with an increasing number of trials (Experiment 3), which supports the idea that the current individual subject modeling could be a viable approach for future clinical research into biomarkers for mental health conditions. Collectively, our findings suggest that the relationship between the velocity of tactile stimulation and subjective pleasantness follows a negative quadratic function in most people, that the shape of this function was robust and replicable across trials and sessions and present for both hairy and non-hairy skin, and thus it is likely to represent a universal function for tactile pleasantness experience in humans.

### The perception of affective touch at the individual level

We believe that our results provide an important contribution to the affective touch field from the experimental, behavioral, and clinical points of view. Several studies have shown that slow, caress-like touch delivered at velocities between 1 and 10 cm/s might optimally activate the CT afferent system and is perceived as more pleasant than slower or faster velocities based on subjective ratings (e.g., Löken et al., 2009; Vallbo et al., 1999). At the group level, the averaged pleasantness ratings of touch delivered at velocities in the range between 0.3 and 30 cm/s follow an inverted-U shape curve, with maximum pleasantness between 1 and 10 cm/s. Here, we provide novel evidence that this pattern is also valid at the individual level and on both hairy and non-hairy skin. This is an important advance for the field because it is individuals – not groups – that experience pleasant touch, and the relationships observed at the group level (through data averaging) may not be representative of the specific perception of individual participants. One of the reasons accounting for why we might have found that a higher percentage of participants at the individual level followed the inverted-U shaped pattern compared to the results of one previous study (Croy et al., 2021) is the fact that all our participants were tested under the same conditions with the tactile stimulation always delivered by the same experimenter. In contrast, Croy et al. 2021 pooled data from 5 separate experiments with the common ground that participants were tested on the forearm; this approach might have increased experimental noise and added additional individual variability, over and above that associated with tactile perception per se.

Importantly, by using our modeling approach, we could, for the first time, investigate the relationship between individual traits and characteristics, namely, depression, anxiety, eating disorders symptomatology, and cardiac interoceptive accuracy, and individual differences in the precise shape of the tactile pleasantness-velocity function. We found that individual differences in self-reported depression influenced the steepness of the quadratic curve, at least when touch was delivered on hairy skin. These results are in keeping with recent evidence suggesting that depressive symptomatology can influence how participants perceive and think about tactile social encounters (Triscoli, Croy & Sailer, 2019). Furthermore, these results can be interpreted in the context of a more general anhedonia, which has traditionally been linked to depression and has been found to affect the perception of tactile pleasantness in people with eating disorders (Crucianelli et al., 2021).

Consistent with the idea of affective touch as an interoceptive modality, our results showed that higher cardiac interoceptive accuracy as measured by means of a classic heartbeat counting task (Schandry, 1981) was related to a better fit of the negative quadratic model for the perception of touch on hairy skin (with a similar but non-significant statistical trend observed for non-hairy skin). In this regard, recent studies have reported non-significant relationships between the perception of affective touch and cardiac interoceptive accuracy (Crucianelli et al., 2018; Crucianelli, Enmalm & Ehrsson, 2021); however, interoceptive accuracy was a predictor of tactile pleasantness in a multisensory integration paradigm only when touch was delivered at the borderline velocity for CT optimality (9 cm/s, Crucianelli et al., 2018). Taken together, these findings suggest that the link between cardiac interoception and tactile pleasantness is more complex than originally thought, and modeling approaches might be better suited to describe such relationships.

Overall, our findings suggest that the perception of affective touch follows a relatively consistent pattern at the individual level; nevertheless, this finding should be contextualized in a broader picture that takes into account other individual differences, such as depressive symptomatology and interoceptive abilities. As such, the subjective perception of affective touch reflects the complexity of studying social touch where context and person matter (e.g., Ellingsen, Leknes, Løseth, Wessberg & Olausson, 2016), and it should not be reduced only to bottom-up processing of the signals triggered by the tactile stimulation of the peripheral receptors in the skin; information regarding social and contextual cues modulates the pleasantness experience at central levels of processing.

### Differences in the perception of touch on hairy and non-hairy skin

We found that the pattern of results was similar for touch delivered on hairy (forearm) and non-hairy (palm) skin. These results are surprising in light of classic findings in the field of affective touch (e.g., McGlone et al., 2012) but consistent with a recent meta-analysis that pooled data from 18 studies and reported no systematic difference in pleasantness ratings in response to affective touch across hairy and glabrous skin when considered in group-level analyses (see Cruciani, Zanini, Russo, Boccardi, & Spitoni, 2021). Notably, here, we demonstrated this finding at the individual level for the first time, which is important because it is only at this level that the precise shape of the velocity-pleasantness functions can be modeled and directly compared across skin types. As such, our results are timely and in line with an ongoing shift in the field, suggesting that a more holistic approach might allow us to better capture the perception of tactile pleasantness, where both bottom-up and top-down signals play a role. To the extent that the subjective perception of touch is the result of the activation of mechanoreceptive afferents on the skin in combination with central processing (Ackerley et al., 2014a; McGlone et al., 2014), here, we showed that the pattern of relationship between velocity of touch (and presumably CT activation; Löken et al., 2009) and subjective tactile perception follows a similar pattern across hairy and non-hairy skin at the individual level. In the affective touch field, it has long been common practice to select a glabrous skin site (i.e., palm of the hand) as a control condition to compare to the perception of tactile pleasantness on hairy skin. However, our findings might suggest that more careful consideration is needed when selecting control conditions and in interpreting the results when any remarkable difference is found between hairy and non-hairy skin sites because such findings may not relate to tactile pleasantness per se. In fact, our findings indicate that the inverted-U shaped curve could constitute a fundamental function that describes the typical relationship between tactile stimulation velocity and subjective pleasantness across all body parts and skin types (although further studies are needed to confirm this across more body parts than the three sites investigated here).

Nevertheless, we found that interoceptive accuracy and self-reported depression were related to the perception of tactile pleasantness on hairy skin only, thus indirectly suggesting that there are intrinsic differences in the processing and interpretation of touch based on skin site (e.g., Ackerley et al., 2014a; Kirsch et al., 2018). As such, our results further contribute to our understanding of touch perception and, in particular, the relative contribution of the CT system to the perception of tactile pleasantness. A recent study reported sparse CT afferent recordings from glabrous skin for the first time (Watkins et al., 2021); thus, it could be that even a small number of CT fibers is sufficient to contribute to the perception of tactile pleasantness from a physiological point of view, and therefore, any difference previously observed across skin sites might be due to other factors, such as top-down beliefs about touch perception (e.g., Crucianelli et al., 2021) or previous tactile experiences throughout life (Beltrán, Dijkerman & Keizer, 2020).

### Stability of touch perception across testing sessions and repetitions

The results of Experiment 2 suggest that healthy participants present a rather consistent relationship between velocity of touch and tactile pleasantness at the individual level across two testing sessions. Thus, the way in which we perceive tactile pleasantness might represent a trait rather than a state characteristic. This finding is particular important for clinical research because it can pave the way for further investigation exploring whether the perception of affective touch can be used as a behavioral biomarker for the early identification and diagnosis of psychiatric and neurodevelopmental disorders.

Several studies in the field of affective touch have been criticized for the low number of repetitions used at each velocity. However, it has also been highlighted that several repetitions could reduce the perceived pleasantness of touch (Triscoli, Ackerley, & Sailer, 2014). In this context, the results of Experiment 3 showed that increasing the number of repetitions at each velocity from 3 to 10 did not significantly deteriorate the fit of the model. That is, participants already showed a strong pattern after 3 trials, and this pattern did not change over subsequent trials. These results provide additional, indirect validity for the findings from previous studies that investigated the perception of affective touch using less than 10 repetitions per velocity (see Cruciani, Zanini, Russo, Mirabella, Palamoutsi & Spitoni, 2021 for a recent review), and they further suggest that the perception of affective touch might represent a trait characteristic, with a strong potential to be used as such in future studies.

In this regard, we believe that our results provide important methodological information to consider when designing future studies. The evidence that it is possible to reliably assess affective touch perception using a low number of trials might be crucial for studies conducted with clinical populations presenting symptoms such as fatigue, cognitive impairment, and difficulties in social interactions, as well as when testing in hospital facilities or at bedsides. In such situations, it is essential to collect reliable data while keeping the testing time as brief as possible to guarantee participants’ comfort and safety.

### Concluding remarks and future directions

We believe that our findings highlight the translational potential of quantifying the perception of affective touch as a proxy for socially embodied cognition and have important implications for our understanding of the relationship between mental health and affective touch (Croy et al., 2016). The affective touch system seems to play a role in the development and maintenance of affiliative behaviors and social bonding and for the communication of emotions (see Olausson et al., 2002; Löken et al., 2009; Kirsch et al., 2018; McGlone et al., 2007; Morrison, Löken & Olausson, 2010). Thus, it has been suggested that this modality could be important for the development of the social brain (see Walker & McGlone, 2013 for a review) and for the way we relate to ourselves and others (Crucianelli et al., 2013). Furthermore, the integration of interoceptive/affective (including signals derived from affective cutaneous stimulation) and exteroceptive/sensory information plays a critical role in bodily awareness at any given time and the construction of the subjective experience of the self (e.g., Critchley et al., 2004; Craig, 2009; Crucianelli et al., 2018; Salvato et al., 2020). Our findings that healthy participants showed a rather consistent relationship between velocity of touch and tactile pleasantness at the individual level, combined with clinical data suggesting a close link between affective touch and mental health, can pave the way for further investigation exploring the perception of affective touch in people at different stages of mental health conditions, ranging from populations at risk of developing such disorders to people who have successfully recovered from them. For example, it remains unclear whether disruptions in the perception of affective touch can be considered a consequence of other symptoms or whether they are a contributing factor for the development of mental health disorders. Along this line, one study investigated the perception of affective touch in people who had recovered from anorexia nervosa compared to women with anorexia nervosa and matched healthy controls (Crucianelli et al., 2021). The results suggested that the difficulties in the perception of affective touch might persist even after an otherwise successful recovery. Thus, such deficits might represent a trait rather than a state phenomenon related to the status of malnutrition associated with this clinical condition.

To the best of our knowledge, this is the first study to apply an advanced modeling approach to better characterize the perception of tactile pleasantness at the individual level. We systematically investigated the stability of the profiles across body sites, comparing hairy and non-hairy skin, across two testing sessions one week apart, and across several repetitions (from 3 to 10). Our results provide an important novel contribution to the field and further validate previous findings investigating the perception of affective touch. Future studies could also include a measure of precision of the subjective perception by asking participants to report their confidence in their responses, for example (e.g., Denison, 2017; Rouault, Seow, Gillan, & Fleming, 2018; Rouault, McWilliams, Allen & Fleming, 2018). Such metacognitive approaches hold the potential to provide a more complete picture of the way peripheral tactile stimulation can give rise to a subjective pleasantness percept.

## Acknowledgments

This work was supported by the Hjärnfonden, the Swedish Research Council (Distinct Professor Grant) and the European Research Council (SELF-UNITY) to Henrik Ehrsson. Laura Crucianelli was supported by the Marie Skłodowska-Curie Individual Fellowship (891175). We would like to thank Adam Enmalm for helping with data collection.

## Author contributions

L.C, M.C. and H.H.E. conceived and designed the experiments. L.C. collected the data. M.C. and L.C. conducted the statistical analysis. L.C., M.C. and H.H.E. wrote the manuscript.

## Materials & Correspondence

Correspondence and material requests should be addressed to.

## Appendix 1

### 1. Method

#### 1.1 Cardiac interoceptive accuracy: heartbeat counting task (HCT)

The experimenter recorded the heartbeat frequency by means of a Biopac MP150 Heart Rate oximeter (Biopac Systems Inc., Goleta, CA, United States) attached to the participant’s non-dominant index finger and connected to a Windows laptop with AcqKnowledge software (version 5.0), which enabled to extract the actual number of heartbeats using the ‘count peak’ function. Attention was given to place the soft oximeter around the finger firmly but not overtightly to reduce the possibility that the participants could perceive heart pulsing from their finger (Crucianelli et al., 2018; Murphy et al., 2019). As part of the task, a 5-minute heartbeat baseline was recorded to check for the presence of autonomic neuropathy. During this time, we presented the instructions for the heartbeat counting task (Schandry, 1981). Participants were asked to silently count their heartbeats between two verbally given signals of ‘go’ and ‘stop’ without manually taking their pulse. Both hands were placed on the table to ensure that no body part was touched. Participants completed a practice trial of 15 seconds before proceeding to the three experimental trials lasting 25 s, 45 s and 65 s, which were presented in a randomized order. No information about the length of the trials was provided. Short breaks of 30 s were given between each trial. By applying a validated formula that allowed us to compare the counted and recorded heartbeats, we obtained a number between 0 and 1, which can be considered a measure of cardiac interoceptive accuracy.

#### 1.2 Precisions regarding residual standard deviations

To compare the relative relevance of each model at the individual level for our whole sample, we performed a confidence interval analysis (CI analysis) on the residual standard deviation (RSD). Simply put, in addition to knowing which model fits best, we wanted to know how much better one model was compared to the other for each participant and whether this difference in performance between the two models was meaningful at the group level. The RSD reflects how much the observed data spread around the regression curve and takes into account the degree of freedom of each model. The smaller the residual standard deviation is, the better the fit and the more predictive the model is.

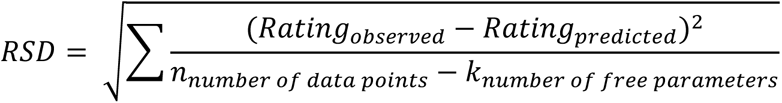

The figure below represents the repartition of the difference between the quadratic and the linear RSD in our population, grouped by profile.

**Appendix Figure 1:**
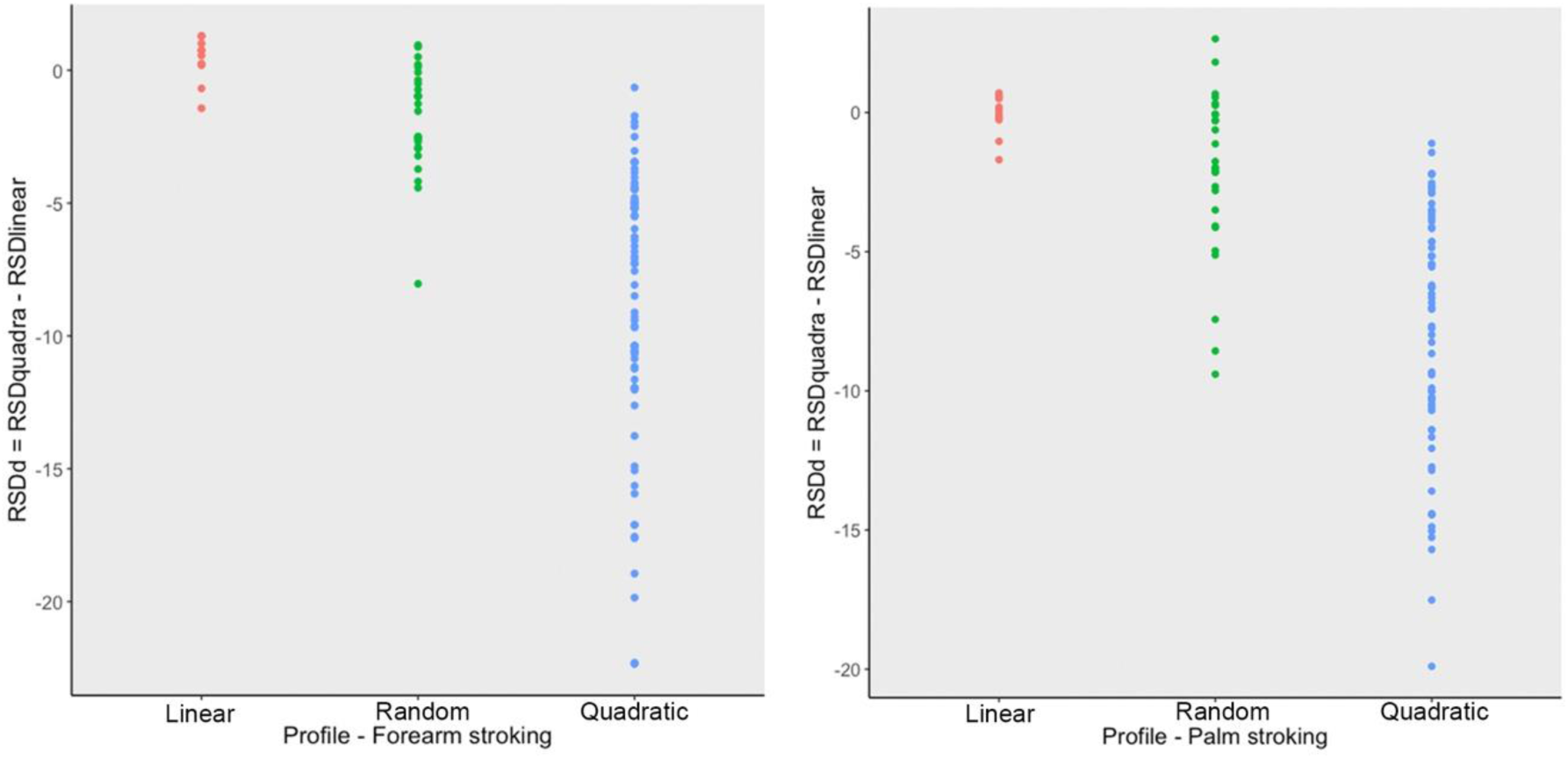
Differences between quadratic and linear RSD (residual standard deviation) plotted by profile. As expected, the RSDd is more negative for the quadratic profile, as the RSD is smaller when the fit is better.

#### 1.3 Precisions regarding the correlation analyses

Using correlation analyses, we wondered whether the individual characteristics we collected could predict how well the participants’ answers would be described by a quadratic model and the coefficients of this quadratic fit. In the appendix figure below, we illustrate how increasing or decreasing one given coefficient (A2, A1, or A0) in the quadratic model influenced the overall predicted response of a participant. We focused our correlation analysis on A2 and A0 because they both reflect one specific aspect of the predicted answer, i.e., the curvature and the amplitude, respectively. We had specific hypotheses regarding the potential link between these aspects of perception and the individual characteristics we collected. However, we did not have a similar strong hypothesis regarding the A1 coefficient, as A1 had a more complex influence on the outcome of the quadratic model.

**Appendix Figure 2:**
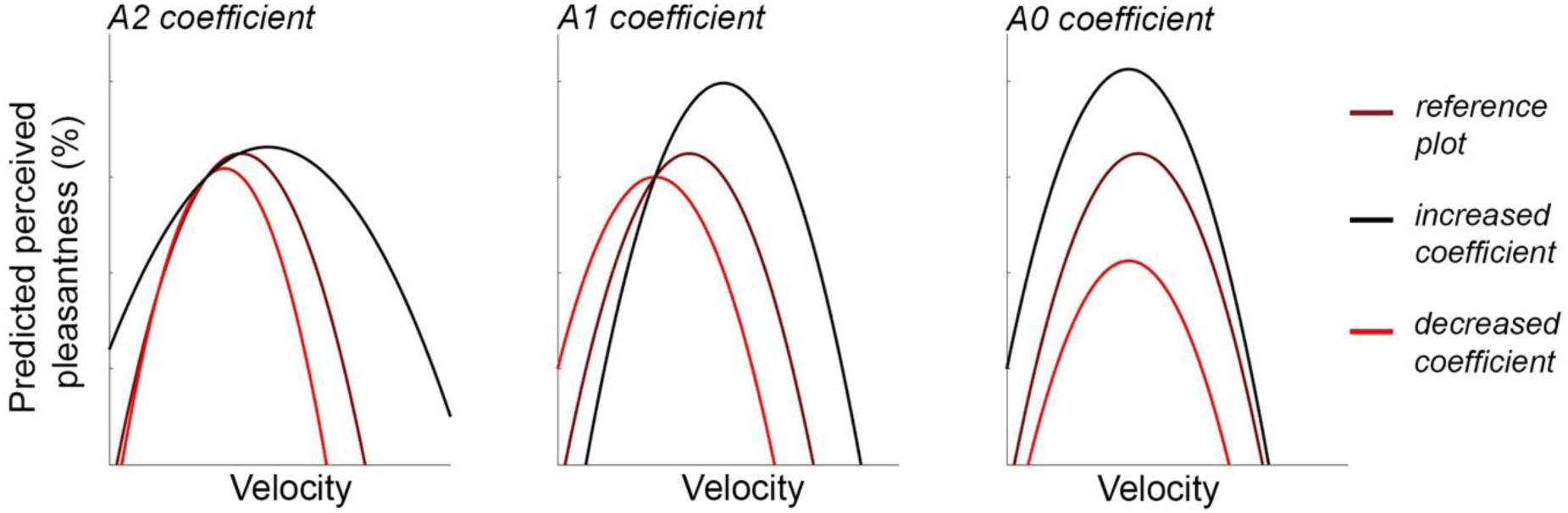
Different quadratic model coefficients differentially impact predicted pleasantness. The quadratic model includes three coefficients: A2 for the quadratic component, A1 for the linear component, and A0 as the intercept. (A2) changes the curvature. (A1) changes the maximal pleasantness rating and the most pleasant velocity. (A0) changes the magnitude of the pleasantness ratings, regardless of the velocity. Brown curve: baseline, red curve: with a decreased coefficient, black curve: with an increased coefficient.

### 2. Results

#### 2.1 Group level results – Individual characteristics

**Appendix Table 1.** Mean and standard deviations for the Eating Disorders Examination Questionnaire (EDE-Q) scores and the depression, anxiety and stress scale scores.

|  | **EDE-Q** | **Depression** | **Anxiety** | **Stress** |
| --- | --- | --- | --- | --- |
| **Total sample** | 1.42 (1.12) | 4.13 (3.87) | 4.02 (3.52) | 5.76 (4.05) |
| **Female (32)** | 1.70 (1.14) | 4.16 (4.05) | 3.38 (2.79) | 5.88 (4.14) |
| **Male (30)** | 1.11 (1.04) | 4.10 (3.74) | 4.70 (4.10) | 5.63 (4.02) |
| **F (p) values** | 4.56 (0.04)* | 0.03 (0.96) | 2.23 (0.14) | 0.05(0.82) |

**Appendix Figure 3:**
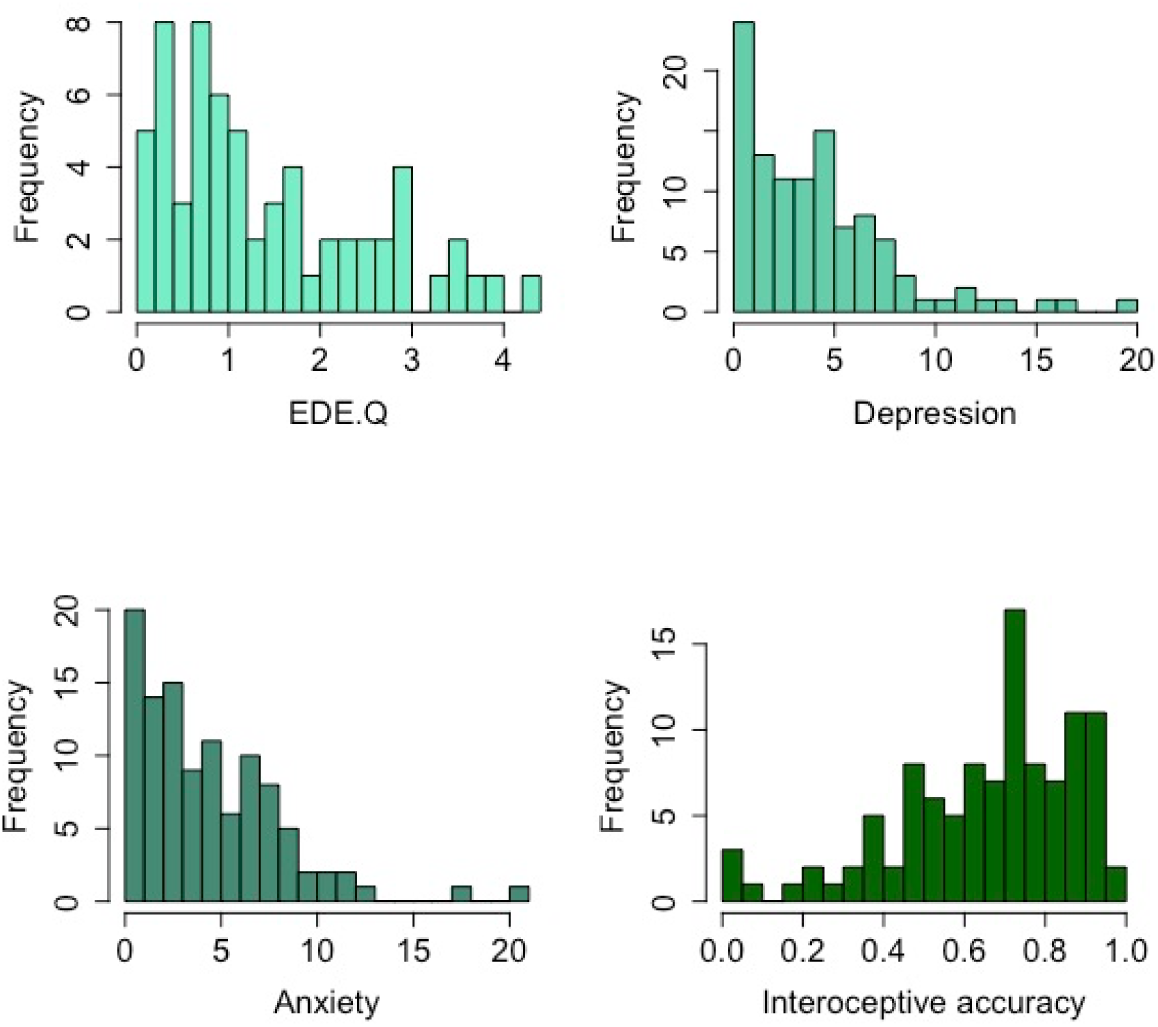
Distribution of the individual characteristic scores among the 107 participants. Participants were asked to complete the following self-report questionnaires: the Eating Disorders Examination Questionnaire (EDE-Q 6.0; Fairburn & Beglin, 1994, 2008; Peterson et al., 2007) and the Depression, Anxiety and Stress Scale–21 Item (DASS, Lovibond and Lovibond, 1995; Henry & Crawford, 2005). Interoceptive accuracy was measured using the heartbeat counting task (Schandry, 1981).

#### 2.2 Group level results - Forearm

The group level results obtained in our experiment were consistent with the patterns observed in most of the studies in the affective touch literature. Below are the group level results for touch applied to the forearm.

**Appendix Figure 4:**
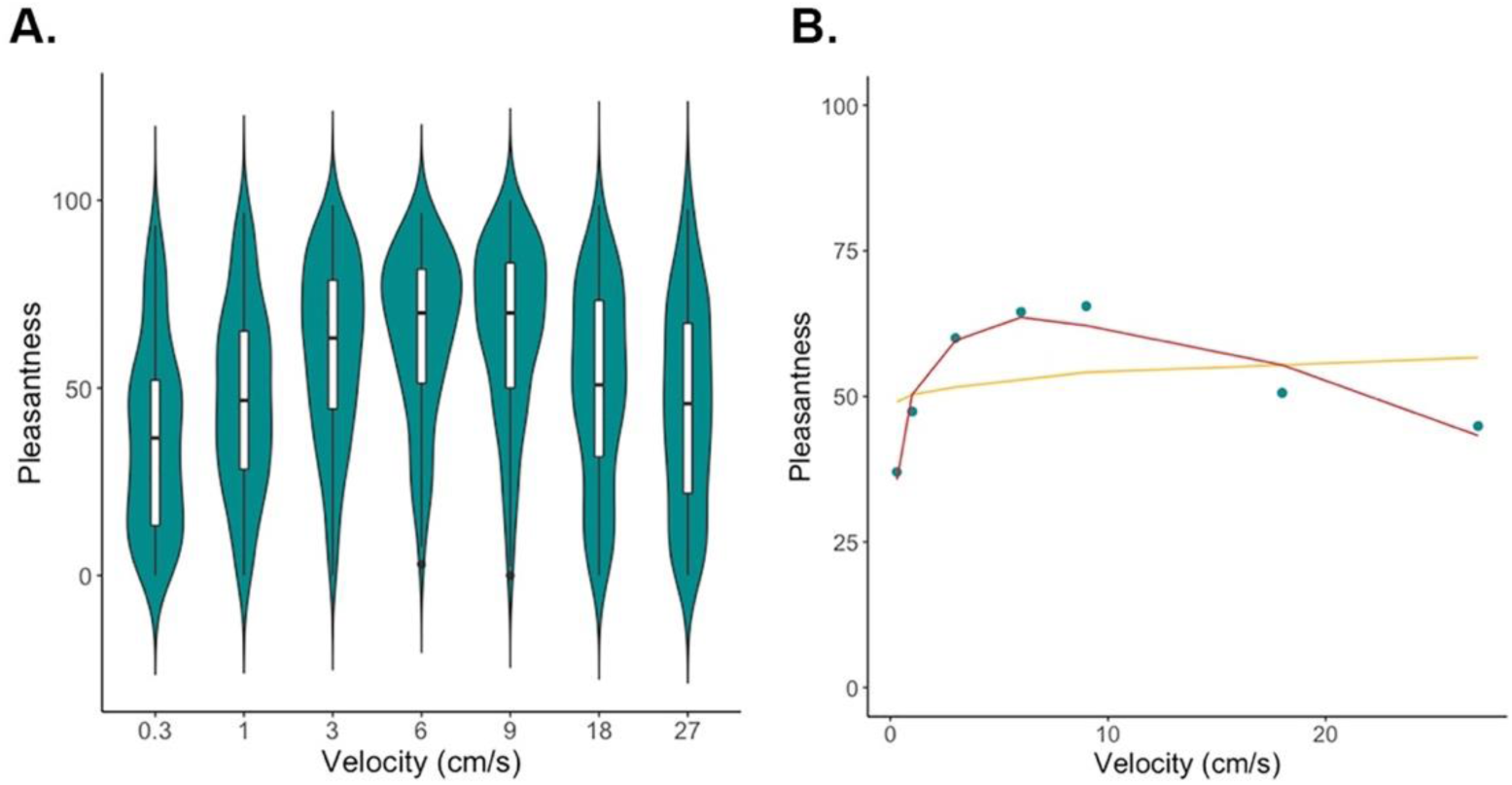
Group level pleasantness ratings in response to strokes applied to the forearm at different velocities. At the group level, we observe the classic inverted U-shaped curve (*F*(2, 4) = 26.93, *p* < 0.005; I(Velocity^2): *t* = -7.078, *p* = 0.00210 **). (A) The plotted colorful shapes reflect the probability densities of the corresponding ratings among the participants. The lower and upper hinges of each box inside the shapes correspond to the first and third quartiles, respectively, with the thick horizontal lines representing the medians. The upper and lower whiskers extending from the boxes indicate the range of maximum to minimum values. (B) Quadratic (red) and linear (yellow) fits of the data (blue dots).

#### 2.3 Fitting procedure: Group level outcome – Forearm

The following plots display the different parameters obtained after the fitting procedure, i.e., the parameter reflecting the goodness of fit of the quadratic model (RMSE) and the relative goodness of fit of the quadratic model compared to the linear model (RSDd), as well as the quadratic model coefficients (A2, A1, A0).

**Appendix Figure 5:**
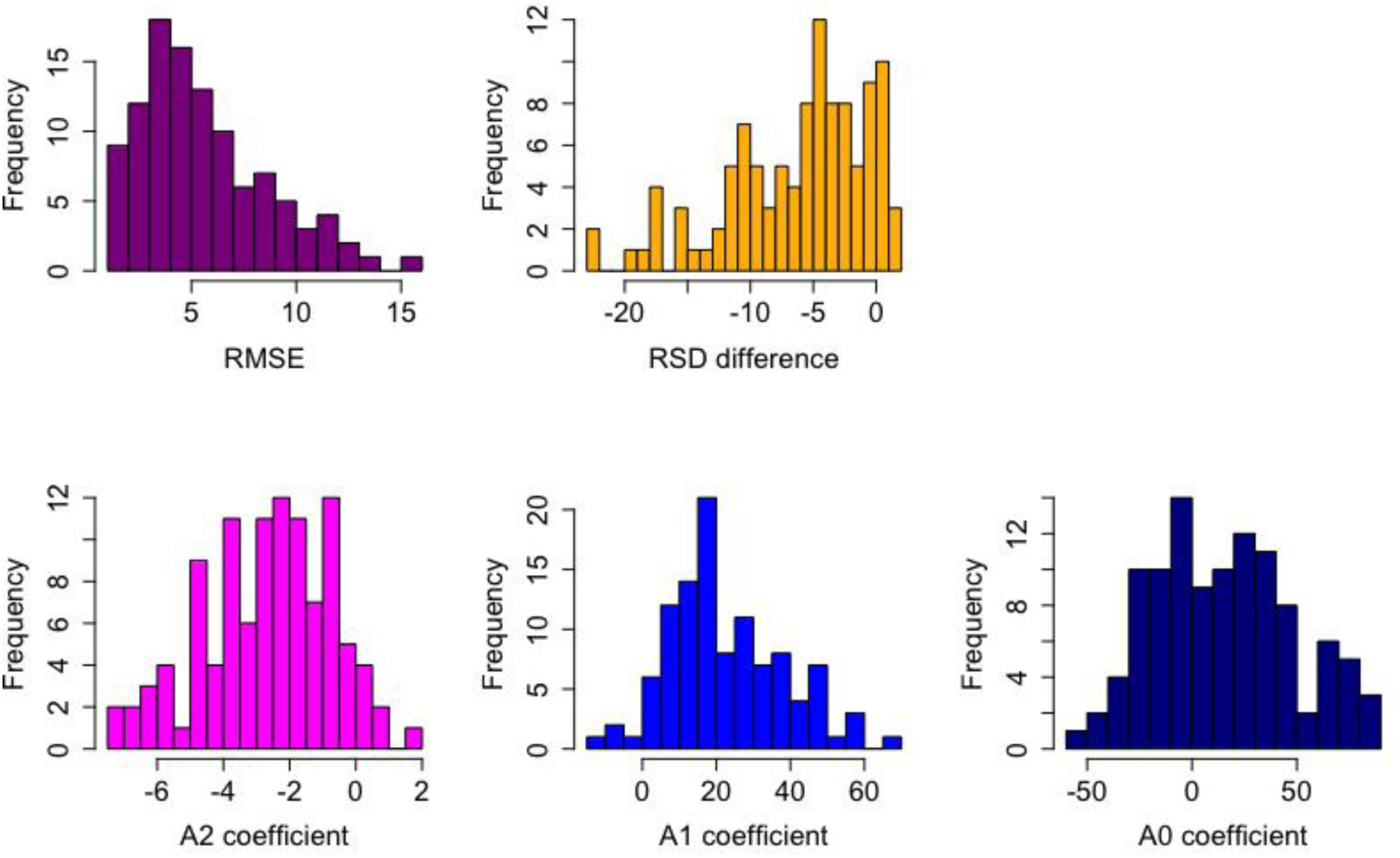
Distributions of the fitting procedure outcomes for Experiment 1.

#### 2.4 Group level results – Palm

The group level results obtained in our experiment were consistent with the patterns observed in most of the studies in the affective touch literature. Below are the group level results for touch applied to the palm.

**Appendix Figure 6:**
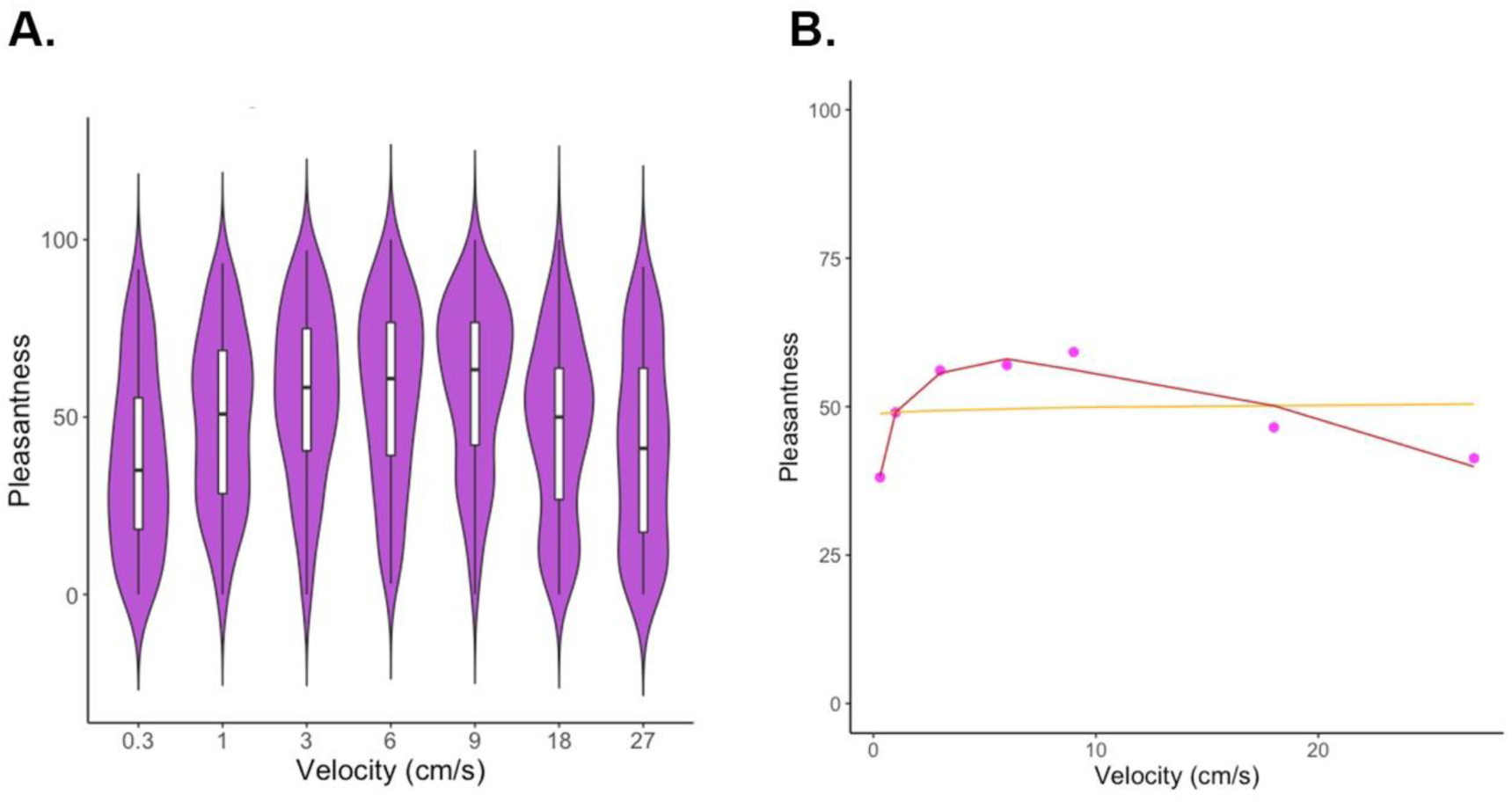
Group level pleasantness ratings in response to strokes applied to the palm at different velocities. At the group level, we observe the classic inverted U-shaped curve (*F*(2, 4) = 29.25, *p* < 0.005, I(Velocity^2): *t* = -7.627, *p* = 0.00159 **). (A) The plotted colorful shapes reflect the probability densities of the corresponding ratings among the participants. The lower and upper hinges of each box inside the shapes correspond to the first and third quartiles, respectively, with the thick horizontal lines representing the medians. The upper and lower whiskers extending from the boxes indicate the range of maximum to minimum values. (B) Quadratic (red) and linear (yellow) fits of the data (purple dots).

#### 2.5 Fitting procedure: Group level outcome - Palm

The following plots display the different parameters obtained after the fitting procedure, i.e., the parameter reflecting the goodness of fit of the quadratic model (RMSE) and the relative goodness of fit of the quadratic model compared to the linear model (RSDd), as well as the quadratic model coefficients (A2, A1, A0).

**Appendix Figure 7:**
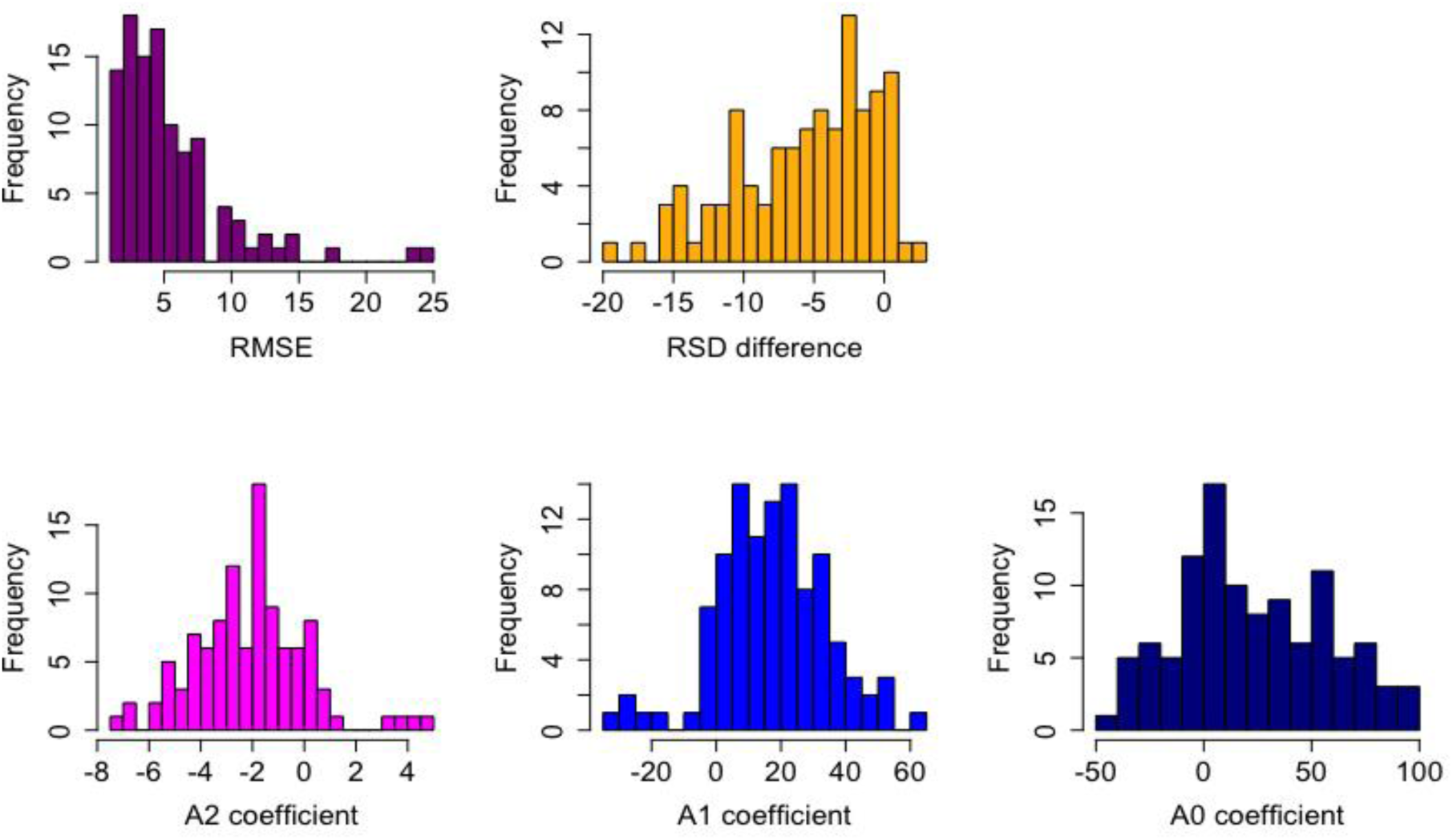
Distributions of the fitting procedure outcomes for Experiment 1.

#### 2.6 Stability of the model’s coefficients for the participants with a ‘quadratic’ profile

This section examines the correlations between the model coefficients A2 (quadratic term), A1 (linear term), and A0 (intercept) for the different body sites for the participants who had a ‘quadratic’ profile with both the forearm and the palm. Each coefficient was significantly correlated with its homolog from the other body site (Figure S8). These strong observed correlations argue in favor of a stability of pleasantness judgment across body sites.

**Appendix Figure 8:**
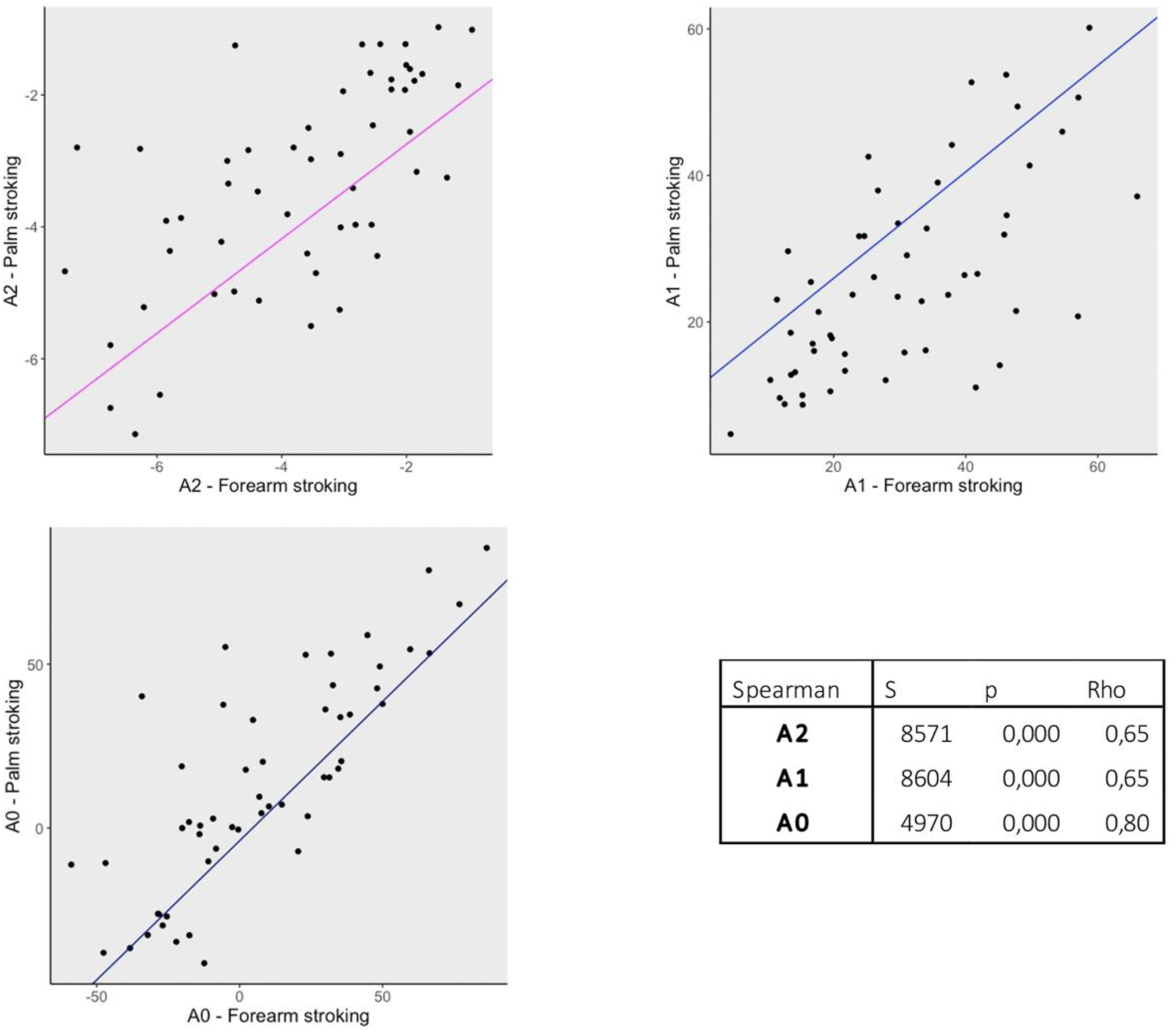
Quadratic model coefficients for participants who had a ‘quadratic’ profile for forearm and palm stimulation in Experiment 1. For the 54 participants who had a ‘quadratic’ profile at both body sites, the quadratic coefficient A2 (A), the linear coefficient A1 (B), and the intercept A0 (C) estimated for the forearm were correlated with those estimated for palm stimulation.

## Appendix 2

### 1. Results

#### 1.1 Group level results - Forearm

**Appendix Figure 9:**
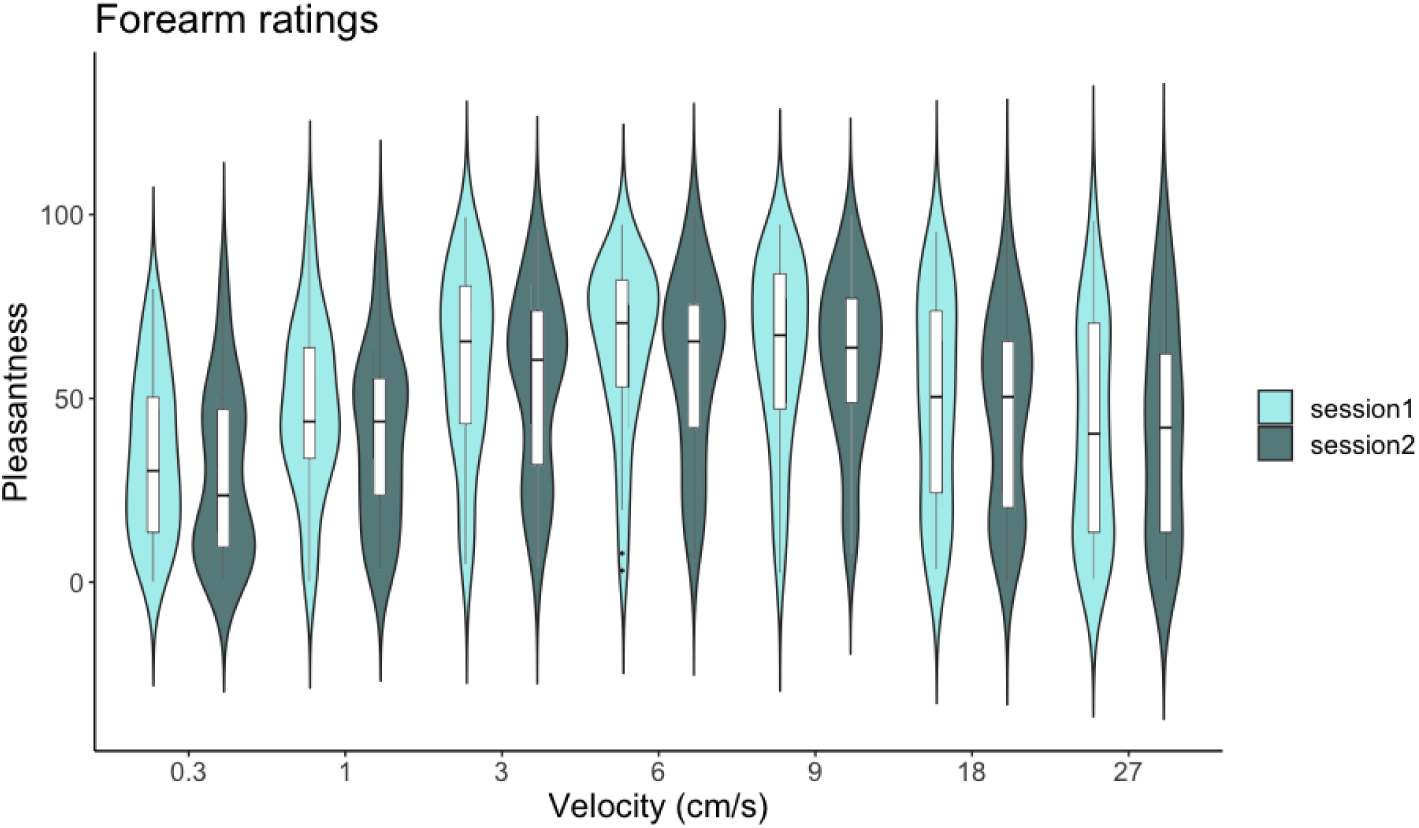
Group level pleasantness ratings in response to strokes applied to the forearm at different velocities in session 1 (light blue) and session 2 (darker blue) (Experiment 2). The plotted colorful shapes reflect the probability densities of the corresponding ratings among the participants.

#### 1.2 Stability across sessions for forearm stroking – ‘quadratic’ profile

We looked at the correlation between the model coefficients A2 (quadratic term), A1 (linear term), and A0 (intercept) between the two sessions for the 25 participants who had a ‘quadratic’ profile in both sessions. Each coefficient was significantly correlated with its homolog from the other session (Table S2 and Figure S10). These strong observed correlations argue in favor of a stability of pleasantness judgment across sessions.

**Appendix Table 2 and Figure 10:**
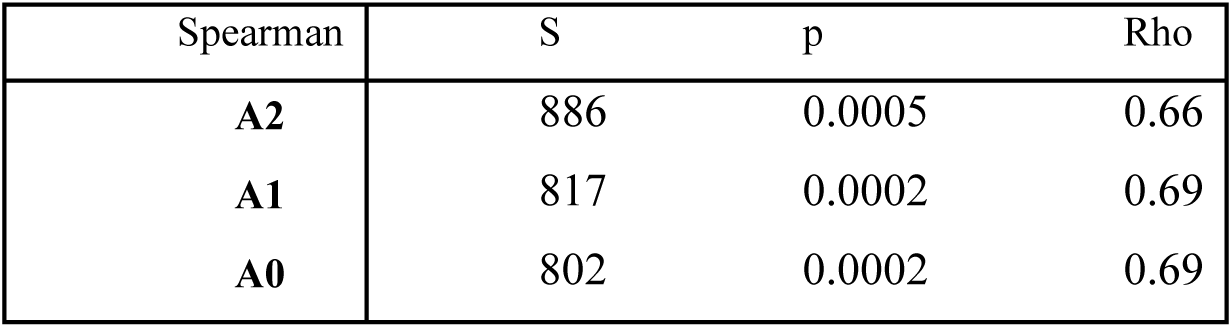
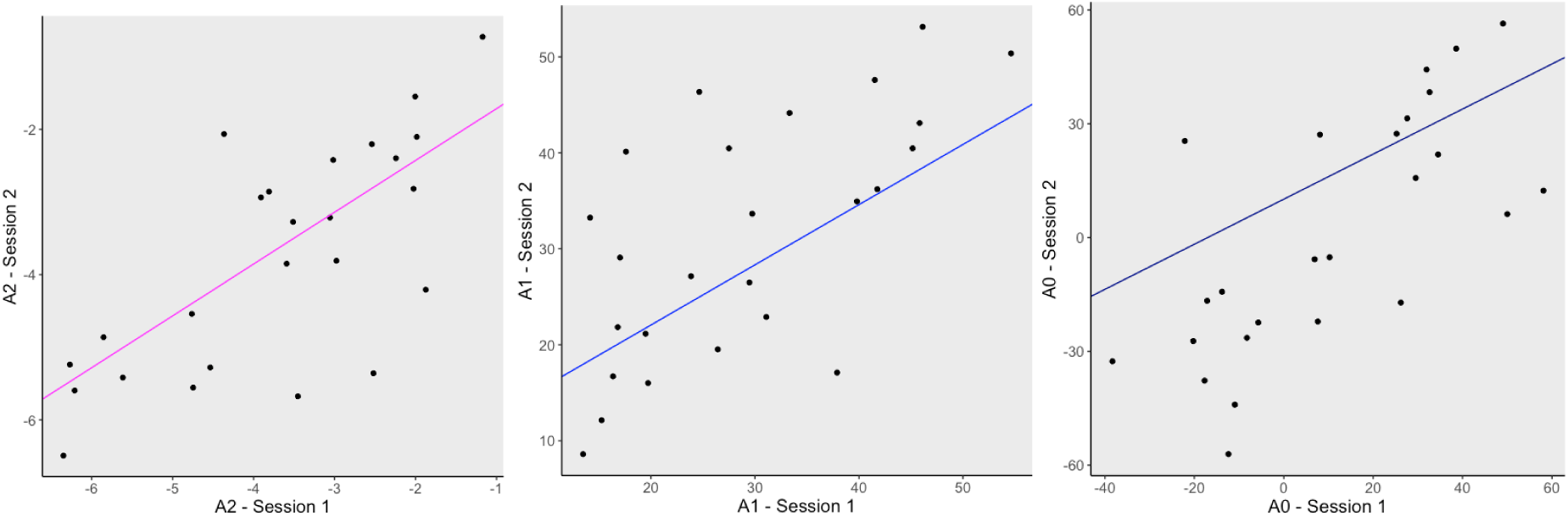
Quadratic model coefficients for participants who had a ‘quadratic’ profile in sessions 1 and 2 (Experiment 2–forearm stimulation). For the 25 participants who had a ‘quadratic’ profile in both sessions, the quadratic coefficient A2 (A), the linear coefficient A1 (B), and the intercept A0 (C) estimated in session 1 are correlated to those estimated in session 2.

#### 1.3 Group level results - Palm

**Figure S11:**
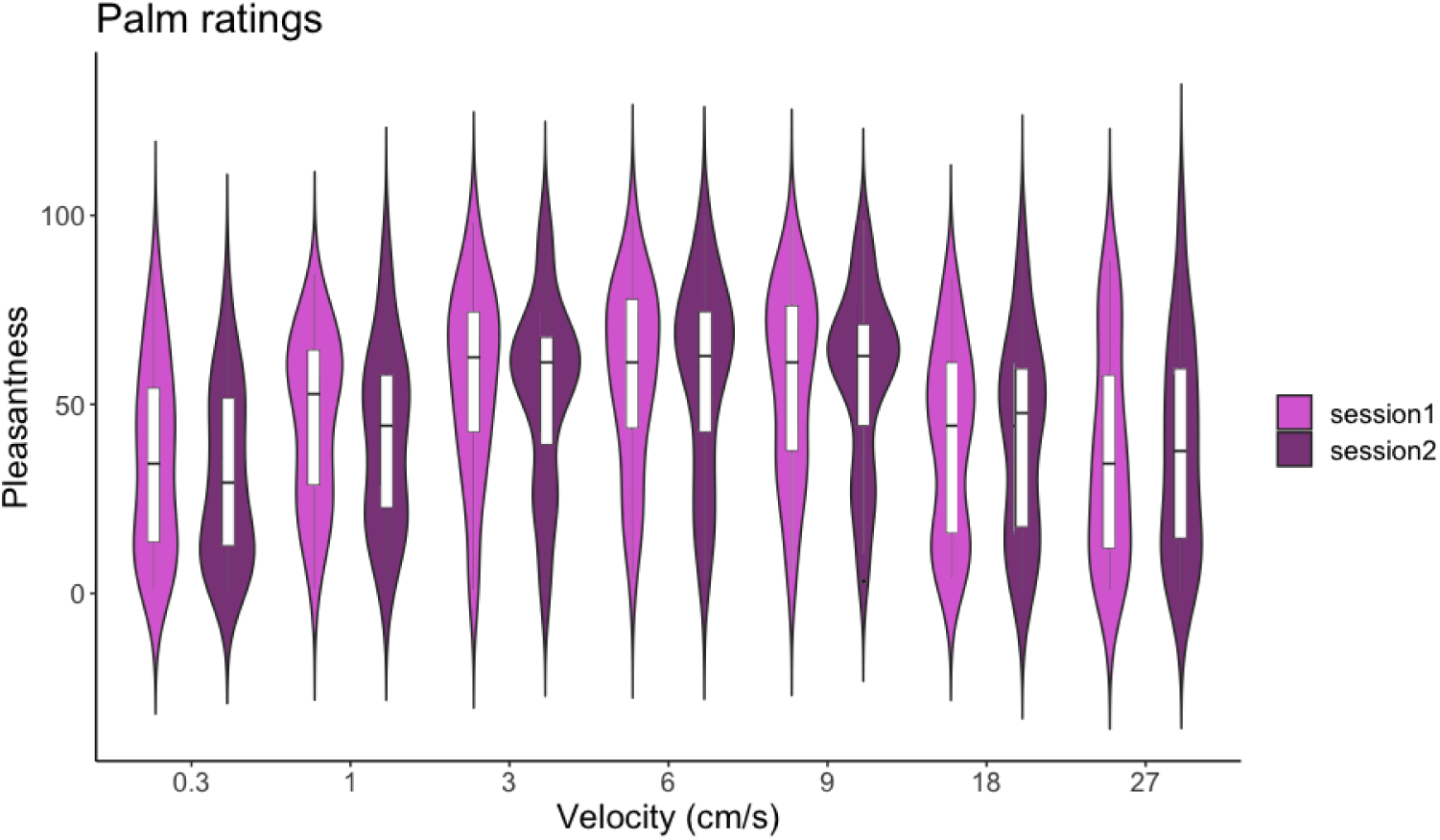
Group level pleasantness ratings in response to strokes applied to the forearm at different velocities in session 1 (light purple) and session 2 (darker purple) (Experiment 2). The plotted colorful shapes reflect the probability densities of the corresponding ratings among the participants.

#### 1.4 Stability across sessions for palm stroking – ‘quadratic’ profile

We looked at the correlation between the model coefficients A2 (quadratic term), A1 (linear term), and A0 (intercept) between the two sessions for the 27 participants who had a ‘quadratic’ profile in both sessions. Each coefficient was significantly correlated with its homolog from the other session (Table S3 and Figure S12). These observed correlations argue in favor of a stability of pleasantness judgment across sessions. However, the correlation between A1 in session 1 and session 2 was weaker than the correlations previously reported.

**Appendix Table 3 and Figure 12:**
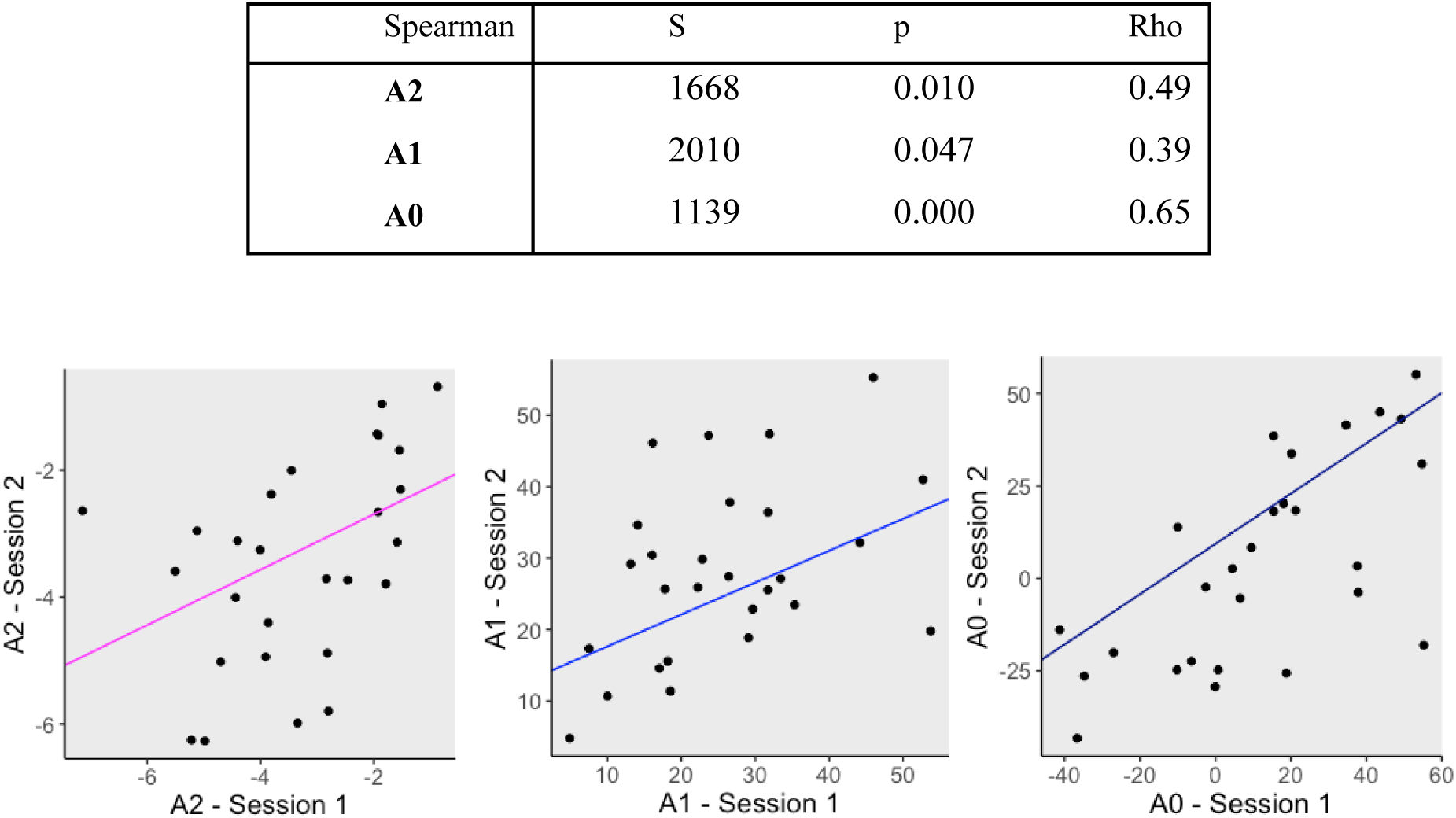
Quadratic model coefficients for participants who had a ‘quadratic’ profile in sessions 1 and 2 (Experiment 2–palm stimulation). For the 25 participants who had a ‘quadratic’ profile in both sessions, the quadratic coefficient A2 (A), the linear coefficient A1 (B), and the intercept A0 (C) estimated in session 1 are correlated to those estimated in session 2.

| Spearman | S | p | Rho |
| --- | --- | --- | --- |
| A2 | 1668 | 0.010 | 0.49 |
| A1 | 2010 | 0.047 | 0.39 |
| A0 | 1139 | 0.000 | 0.65 |

#### 1.5 Group level results – Dorsal hand

##### Response to dorsal hand (hairy skin) stimulation

###### a. Quadratic, linear, and random profile categorization

At the group level, we observed the classic inverted U-shaped curve (Figure S13).

**Appendix Figure 13:**
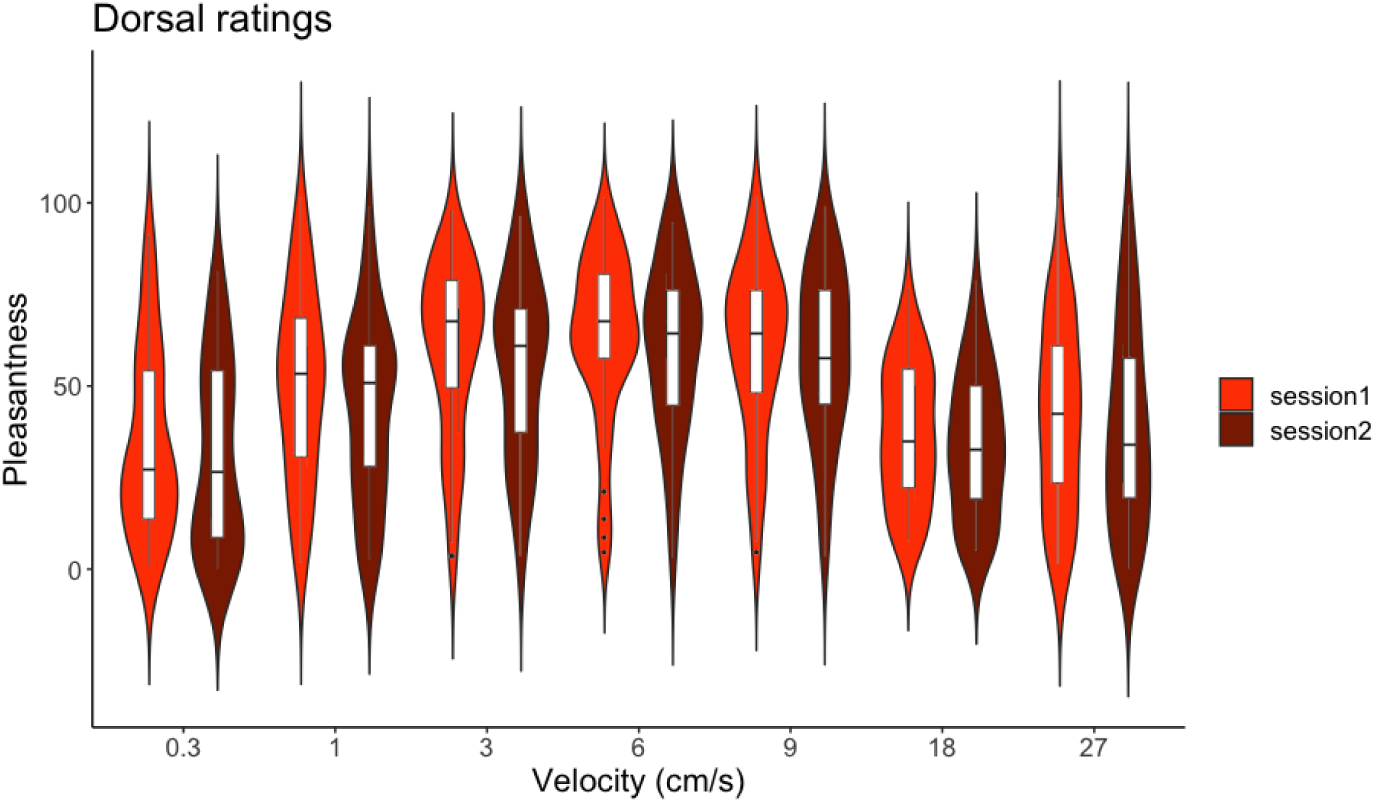
Group level pleasantness ratings in response to strokes applied to the forearm at different velocities in session 1 (light orange) and session 2 (brown) (Experiment 2). The plotted colorful shapes reflect the probability densities of the corresponding ratings among the participants.

In response to stimulation applied to the dorsal hand, 20 (48.8%) participants did not show a significant effect of velocity on perceived pleasantness and were thus categorized as ‘random’, 20 (48.8%) participants were categorized as having a ‘quadratic’ profile, and 1 (2.4%) participant was characterized as having a ‘linear’ profile in session 1 (Figure S14). In session 2, 13 (31.7%) participants were categorized as having a ‘random’ profile, 24 (58.5%) participants were categorized as having a ‘quadratic’ profile, and 4 (9.7%) participants were categorized as having a ‘linear’ profile (Figure S14). For the other body sites, the CI analysis showed a clear superiority of the quadratic model compared to the linear model for both sessions (session 1: lower bound: -351, raw sum: -274, upper bound: -204; session 2: lower bound: -308, raw sum: -248, upper bound: -190). However, for this body site, we observed an increase in participants categorized as ‘random’, i.e., for whom the velocity did not significantly impact the pleasantness rating.

**Appendix Figure 14:**
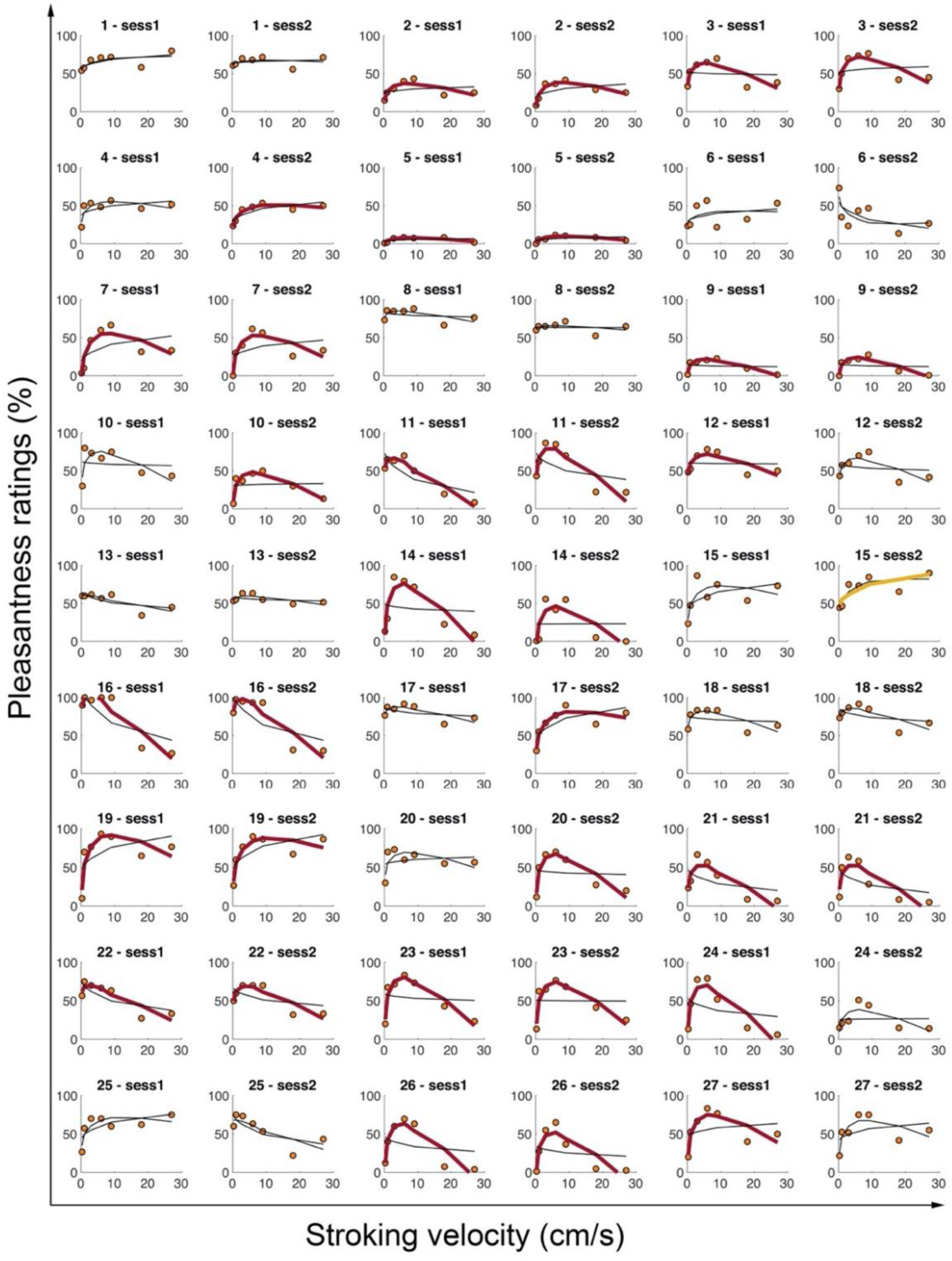
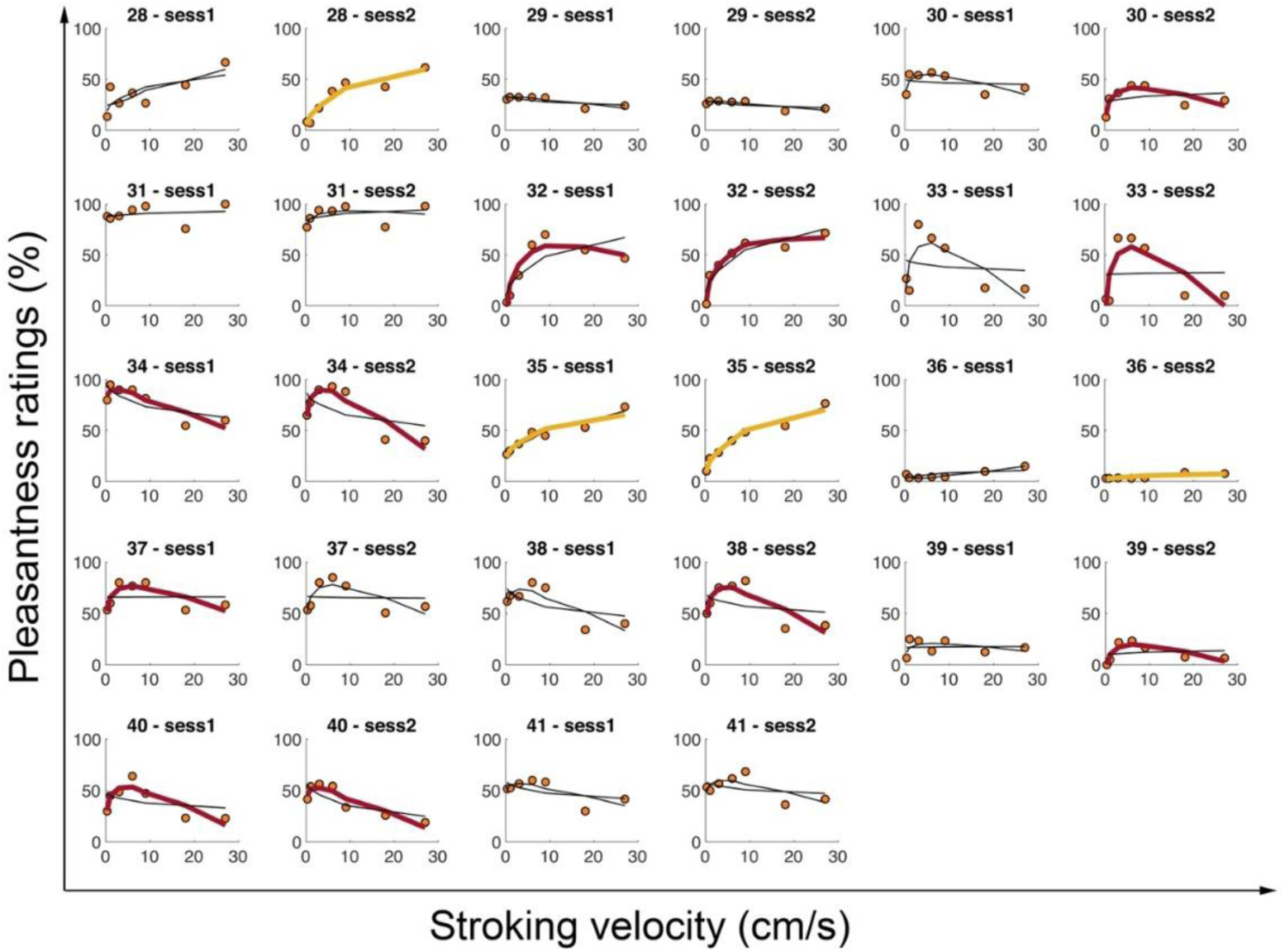
Quadratic and linear fit of individual pleasantness ratings in response to stroking applied at different velocities to the back of the hand in both sessions (Experiment 2). The figure displays the individual pleasantness ratings at each stroking velocity (orange dots) and the linear and quadratic fit results (curves). A thick red curve indicates that the participant has a ’quadratic’ profile (i.e., significant quadratic fit and *p values*(LRT) < .05). A thick yellow curve indicates that the participant has a ‘linear’ profile (i.e., significant linear fit and *p values*(LRT) > .05). The absence of a bold curve (neither red nor yellow) indicates that the participant has a ‘random’ profile. For each participant, the results for both sessions are plotted side-by-side.

###### b. Stability across sessions for the dorsal hand – ‘quadratic profile’

Twenty-six (63%) participants displayed the same profile in session 1 and session 2. This result was above chance level (33%). The detailed repartition of the participants is shown in Table S4. Furthermore, the observed likelihood for each profile exceeded the predicted likelihood for the ‘quadratic’ and ‘linear’ profiles but not the random category (Table S5). This result suggested that the identified profiles were not completely stable across sessions.

**Appendix Table 4:** Repartition of the participants in profiles for both sessions.

|  |  | Session 2 profiles |  |  |  |
| --- | --- | --- | --- | --- | --- |
|  |  | Quadratic | Linear | None | Total |
| Session 1<br>profiles | Quadratic | 16 | 0 | 4 | 20 |
|  | Linear | 0 | 1 | 0 | 1 |
|  | Random | 8 | 3 | 9 | 20 |
|  | Total | 24 | 4 | 13 | 41 |

**Appendix Table 5:** Predicted and observed likelihood for each profile to be identical for both sessions.

|  | Predicted<br>likelihood | Observed<br>likelihood | Difference | Interpretation |
| --- | --- | --- | --- | --- |
| Quadratic | 0.24 | 0.39 | 0.15 | The observed likelihood is higher than the predicted one for the quadratic and linear profile, not for the random profile. |
| Linear | 0.00 | 0.02 | 0.02 |  |
| Random | 0.24 | 0.22 | -0.02 |  |

We then looked at the correlation between the model coefficients A2 (quadratic term), A1 (linear term), and A0 (intercept) between the two sessions for the 16 participants who had a ‘quadratic’ profile in both sessions. Each coefficient was significantly correlated with its homolog from the other session (A2: S = 173, *p* < .001, rho = 0.75; A1: S = 158, *p* < .001, rho = 0.77; A0: S = 86, *p* < .001, rho = 0.87; see Table S6 and Figure S15). These observed correlations argue in favor of a stability of pleasantness judgment across sessions.

**Appendix Table 6 and Figure 15:**
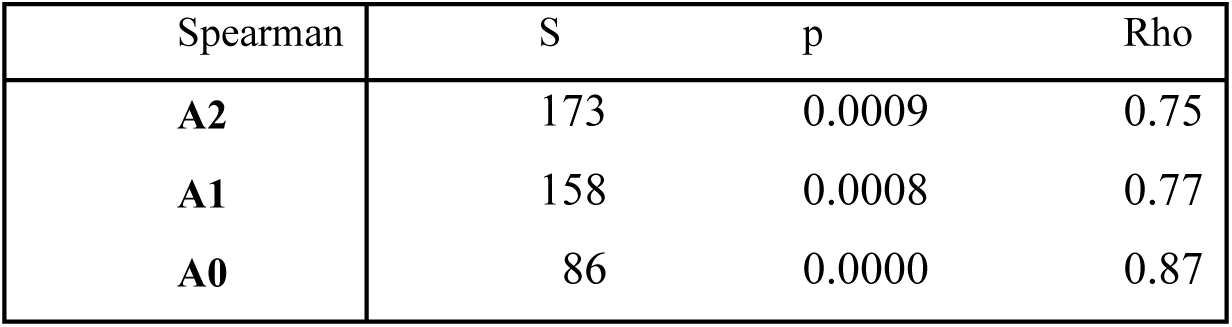
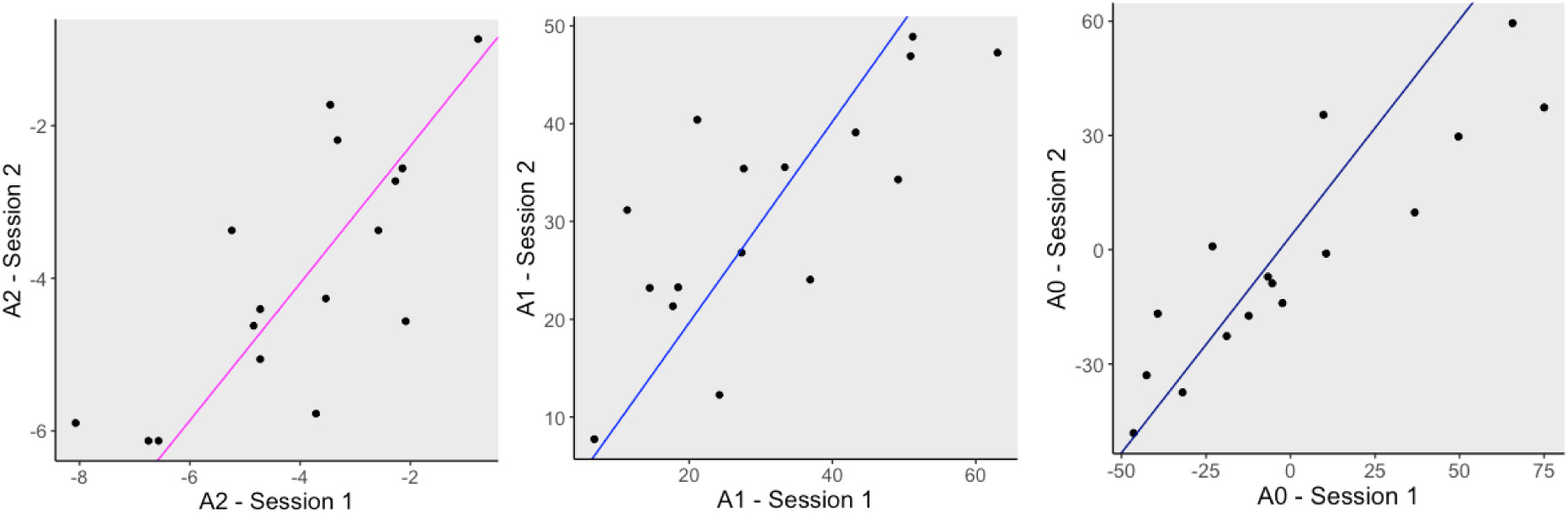
Quadratic model coefficients for participants who had a ‘quadratic’ profile in sessions 1 and 2 (Experiment 2–palm stimulation). For the 25 participants who had a ‘quadratic’ profile in both sessions, the quadratic coefficient A2 (A), the linear coefficient A1 (B), and the intercept A0 (C) estimated in session 1 are correlated to those estimated in session 2.

#### 1.7 Profile repartition across body sites for the subset of participants also tested on the dorsal hand

**1.7 Profile repartition across body sites for the subset of participants also tested on the dorsal hand.**
|  |  | Dorsal stroking |  |  |  |
| --- | --- | --- | --- | --- | --- |
|  |  | Quadratic | Linear | None | Total |
| Forearm stroking | Quadratic | 19 | 0 | 12 | 31 |
|  | Linear | 0 | 1 | 1 | 2 |
|  | Random | 1 | 0 | 7 | 8 |
|  | Total | 20 | 1 | 20 | 41 |
| Palm stroking | Quadratic | 15 | 0 | 15 | 30 |
|  | Linear | 0 | 1 | 2 | 3 |
|  | Random | 5 | 0 | 3 | 8 |
|  | Total | 20 | 1 | 20 | 41 |

## Appendix 3

### 1. Results

#### 1.1 Group level results – Forearm and Palm

**Appendix Figure 16:**
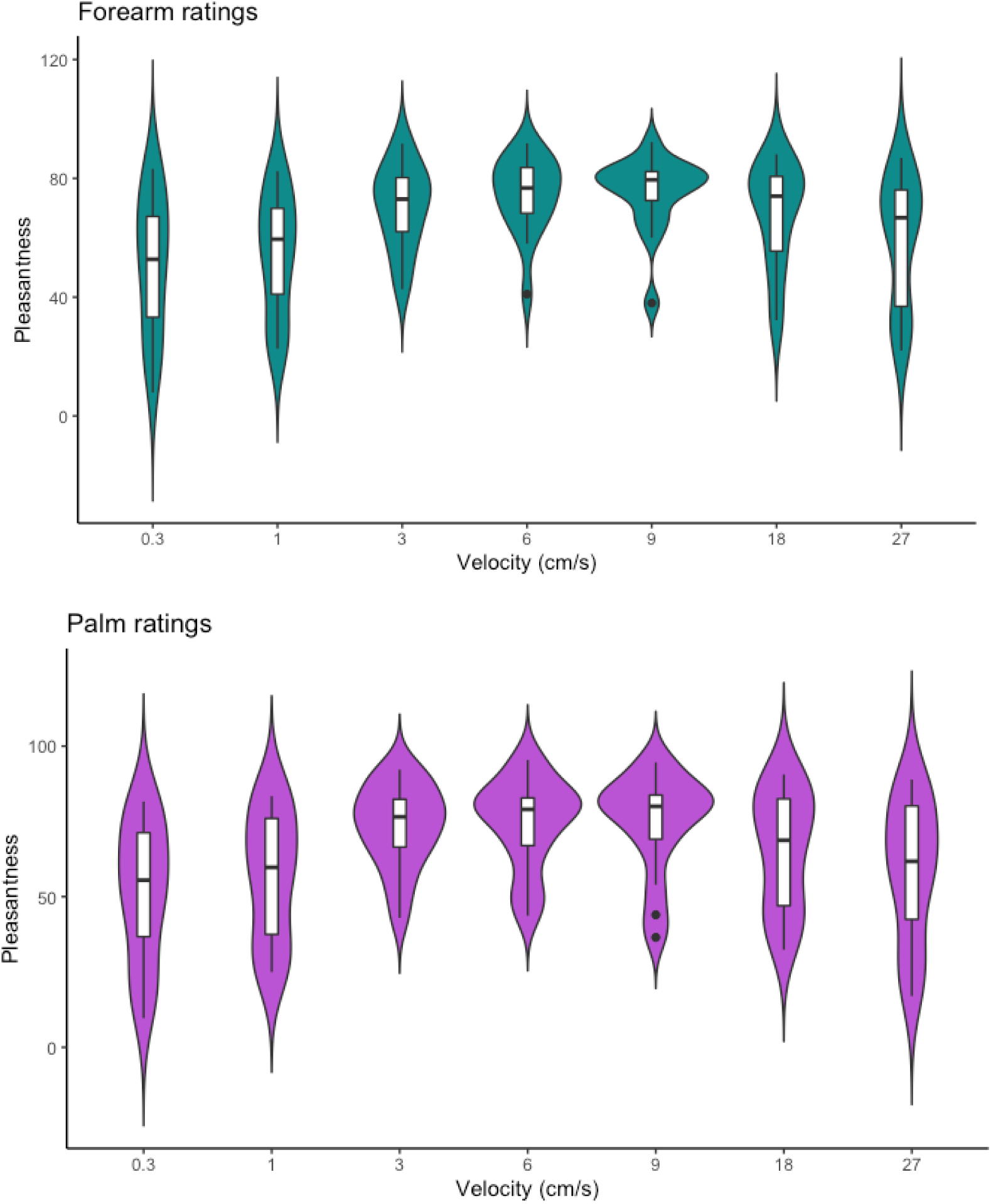
Group level pleasantness ratings in response to strokes applied to the forearm (blue) and the palm (purple) at different velocities (Experiment 3). The plotted colorful shapes reflect the probability densities of the corresponding ratings among the participants.

**Appendix Figure 17:**
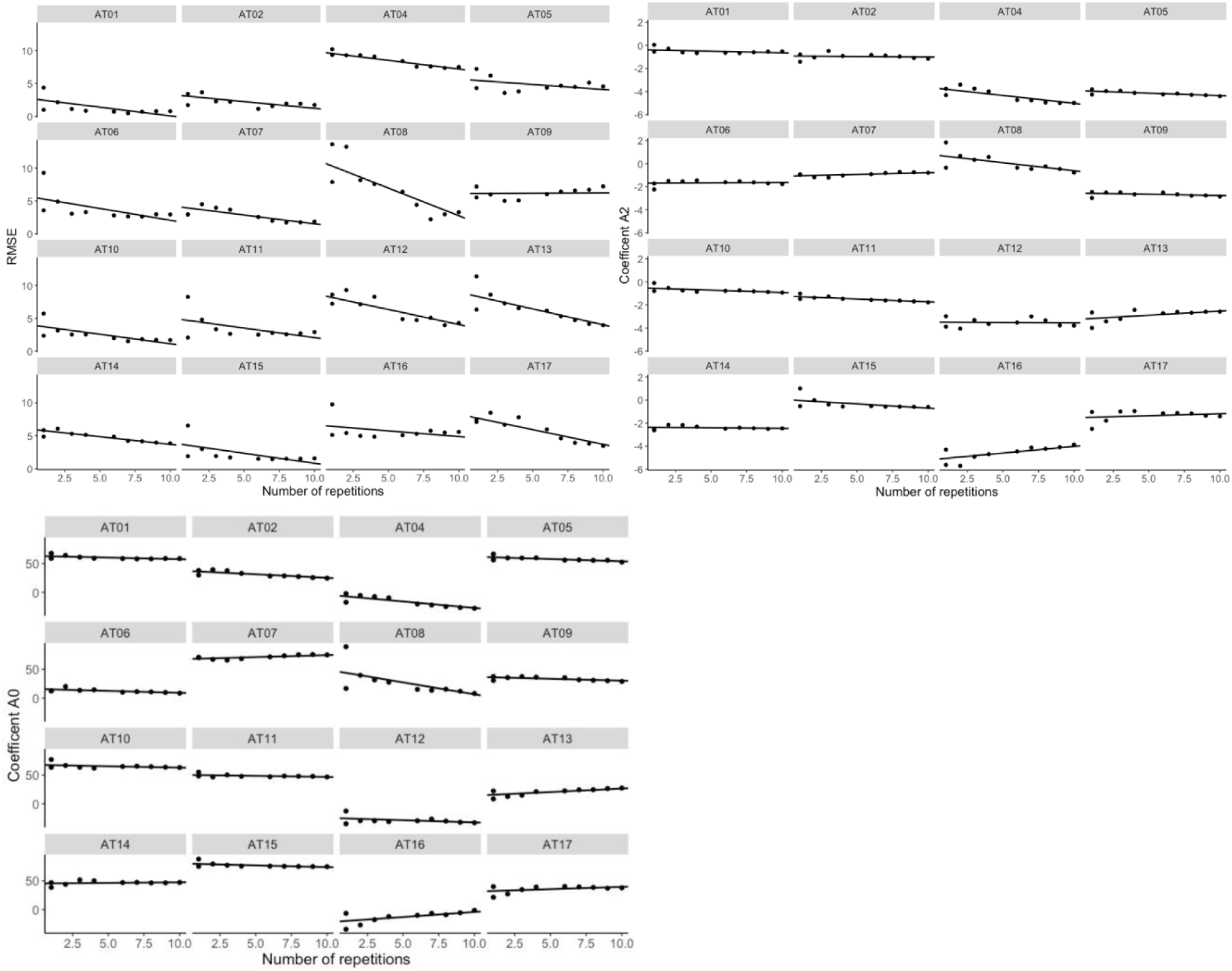
Change in quadratic regression outcomes when the fitting procedure takes into account one to ten repetitions of each stroking velocity on the forearm. A. RMSE. B. A2. C. A0

**Appendix Figure 18:**
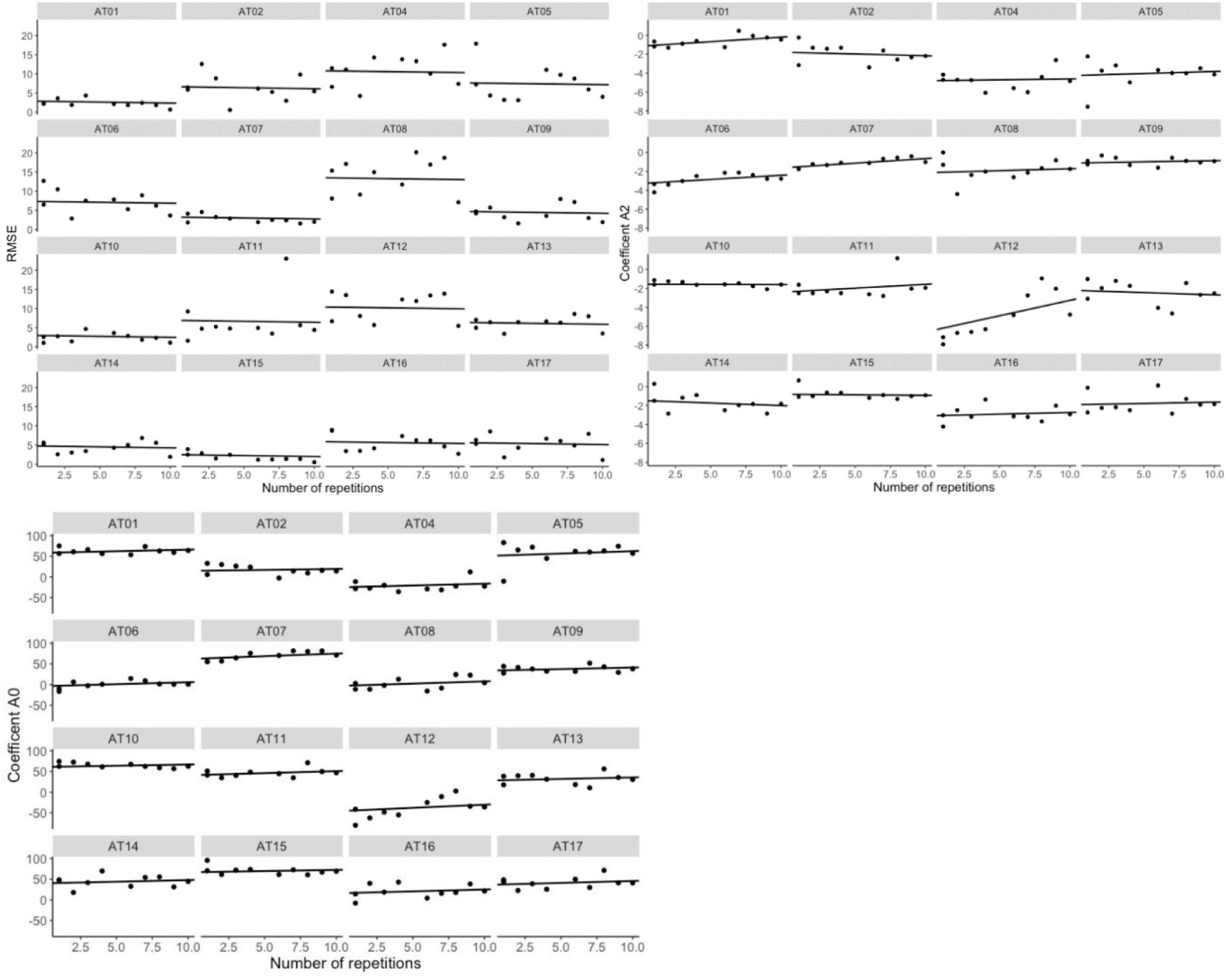
Change in quadratic regression outcomes when the fitting procedure takes into account one to ten repetitions of each stroking velocity on the palm. A. RMSE. B. A2. C. A0

